# Connectome-based Predictive Models of General and Specific Executive Functions

**DOI:** 10.1101/2024.10.21.619468

**Authors:** Shijie Qu, Yueyue Lydia Qu, Kwangsun Yoo, Marvin M. Chun

## Abstract

Executive functions, the set of cognitive control processes that facilitate adaptive thoughts and actions, are composed primarily of three distinct yet interrelated cognitive components: Inhibition, Shifting, and Updating. While prior research has examined the nature of different components as well as their inter-relationships, fewer studies examined whole-brain connectivity to predict individual differences for the three cognitive components and associated tasks. Here, using the Connectome-based Predictive Modelling (CPM) approach and open-access data from the Human Connectome Project, we built brain network models to successfully predict individual performance differences on the Flanker task, the Dimensional Change Card Sort task, and the 2-back task, each putatively corresponding to Inhibition, Shifting, and Updating. We focused on grayordinate fMRI data collected during the 2-back tasks after confirming superior predictive performance over resting-state and volumetric data. High cross-task prediction accuracy as well as joint recruitment of canonical networks, such as the frontoparietal and default-mode networks, suggest the existence of a common executive function factor. To investigate the relationships among the three executive function components, we developed new measures to disentangle their shared and unique aspects. Our analysis confirmed that a shared executive function component can be predicted from functional connectivity patterns densely located around the frontoparietal, default-mode and dorsal attention networks. The Updating-specific component showed significant cross-prediction with the general executive function factor, suggesting a relatively stronger role than the other components. In contrast, the Shifting-specific and Inhibition-specific components exhibited lower cross-prediction performance, indicating more distinct and specialized roles. Given the limitation that individual behavioral measures do not purely reflect the intended cognitive constructs, our study demonstrates a novel approach to infer common and specific components of executive function.

## Introduction

Executive function (EF), also referred to as cognitive control or executive control, is the process of aligning one’s cognition and behavior with current goals (Baddeley, 2000; Diamond, 2013; Miller & Cohen, 2001; Seeley et al., 2007). EF may be categorized into three main processes (Miyake et al., 2000; Lehto et al., 2003): Inhibition, Shifting, and Updating. Inhibition involves actively suppressing the dominant response and is assessed through tasks like the Stroop task (Stroop, 1935), the Flanker task (Eriksen & Eriksen, 1974), and the Stop Signal task (Logan & Cowan, 1984). Shifting, sometimes called cognitive flexibility, occurs when individuals switch between multiple tasks, measured through tasks like the Wisconsin Card Sorting Test (Berg, 1948), the Dimensional Change Card Sort (Frye et al., 1995), and the Color/Shape Switching tasks (Hayes et al., 1998). Lastly, Updating involves continually monitoring pertinent information and integrating it into our finite working memory, which can be measured via tasks like the N-Back task (Kirchner, 1958), the Sternberg task (Sternberg, 1966), and the backward Corsi task (Isaacs & Vargha-Khadem, 1989).

The relationship between different forms of EF has been extensively studied in behavioral experiments. Using large batteries of cognitive tasks, Miyake and colleagues (2000) originally discovered that the performances of all tasks can be loaded onto the three factors: Inhibition, Shifting, and Updating. However, these three factors are intercorrelated. In an attempt to refine the model, Friedman and Miyake (2017) replaced the original Inhibition factor with a general EF factor that is highly loaded by all tasks, resulting in a model with a better fit and more orthogonal factors, which they termed as the “Unity and Diversity” of EF (Friedman & Miyake, 2017). The different EFs are interdependent. For instance, people with superior working memory performance are also more likely to achieve a higher Inhibition score, suggesting that successful working memory may rely on actively inhibiting distractors irrelevant to one’s goals (Conway et al., 2001; Mj & Rw, 2003; Unsworth et al., 2004; Kane et al., 2007). Similarly, Shifting also displays a positive relationship with working memory (Baddeley et al., 2001; Emerson & Miyake, 2003) and Inhibition (Mayr & Keele, 2000; Koch et al., 2010).

Beyond the behavioral level, researchers have long strived to characterize the neural systems underlying each EF component. Earlier lesion studies indicated that damage to the prefrontal cortex may impair performance in different EF tasks, but the exact anatomical locations differed across studies and the type of process targeted (Aron et al., 2003, 2004; Floden & Stuss, 2006; Barbey et al., 2013). Empirical studies and meta-analyses of fMRI data have also revealed multiple brain regions such as the lateral prefrontal cortex, posterior parietal cortex, and dorsal anterior cingulate cortex with both similar and different roles across various EF factors (Assem et al., 2024; Derrfuss et al., 2005; Jamadar et al., 2010; Kim et al., 2012; McNab et al., 2008; Niendam et al., 2012; Rodríguez-Nieto et al., 2022). Discrepant results across studies may lead to different conclusions being drawn about the relationship between EF components, such as the superordinate role of the Updating component (Lemire-Rodger et al., 2019; Rodríguez-Nieto et al., 2022) or the existence of a core cognitive control network in the brain (Niendam et al., 2012).

While univariate fMRI analyses inform us about the brain regions whose activity is modulated by task conditions, connectivity-based approaches enable us to peek into the interaction between different brain regions and how they may be related to individual differences in task performance (Friedman & Robbins, 2022a; Menon & D’Esposito, 2022). In the realm of EF, researchers have shown that the prefrontal cortex’s global functional connectivity as well as structural connectivity profiles can predict one’s EF performance (Cole et al., 2012; Smolker et al., 2015). Furthermore, performance in the three types of EF tasks may correspond with different sets of functional connectivity profiles: Inhibition was highly related to the connectivity strength between the frontoparietal and cingulo-opercular networks (Deck et al., 2023); Shifting scores were correlated with the strength of connectivity within the cingulo-opercular network (Reineberg & Banich, 2016), between the default mode and the dorsal attention networks (Deck et al., 2023), and between the medial frontal and default mode networks (Chén et al., 2019). Updating performance was reflected in the frontoparietal network connectivity profiles (Reineberg & Banich, 2016), and between the angular gyrus and ventral attention network (Reineberg et al., 2015). Another insightful study found that connectivity between the cerebellum and the frontoparietal network predicts a general EF score (Reineberg et al., 2015), but their analysis was limited to resting-state data, and the use of an ICA approach restricted their focus to highly synchronized brain regions.

Leveraging the whole-brain connectivity features, the Connectome-based Predictive Modeling (CPM) method (Finn et al., 2015; Shen et al., 2017) has become a mainstream method for predicting individual differences in cognition. CPM has previously shown exceptional capability in predicting individual cognitive performance such as sustained attention (Rosenberg et al., 2016; Yoo, Rosenberg, Kwon, Lin, et al., 2022), fluid intelligence (Finn et al., 2015; Yoo et al., 2019), creativity (Beaty et al., 2018), and working memory (Avery et al., 2020).

In this study, we extended the application of CPM to study the unity and diversity of the EF task measures of Inhibition, Shifting, and Updating. Going beyond past work, the CPM approach can also predict executive function in novel individuals. For our analyses, while both resting-state and 2-back task data were used to build CPMs, we focused our analyses on using task-based fMRI data over resting-state data for building CPMs due to its higher data quality (Huijbers et al., 2017), greater test-retest reliability (Kristo et al., 2014; Rosazza et al., 2014; Wang et al., 2017), and superior predictive accuracy (Greene et al., 2018, 2020; Yoo et al., 2018).

The behavioral scores and neuroimaging data used in this project were obtained from the open-access Human Connectome Project (HCP) dataset (Van Essen et al., 2012). We used their Flanker task as an approximate measure for Inhibition, the Card Sort task for Shifting, and 2-back tasks for Updating. These EF component assignments to tasks are rough proxies, as we assume task impurity, that is, any given task involves multiple EFs and other cognitive processes (Miyake et al., 2000). Indeed, our goal is to use CPM as a novel approach to tease out shared vs. distinct EF components of these behavioral tasks.

Using 2-back CPM, we explored the unity and diversity of EF at the connectome level from two distinct perspectives. First, we developed CPM models based on the three raw EF measures and achieved significant predictive performance. For each task model, we further identified their underlying canonical functional networks. We also evaluated each task model’s cross-prediction performance to the other tasks (not used for training), and this allowed us to explore the separability and interdependence of the underlying EF components. That is, when a given model can also predict individual differences in tasks different from the one used for model training, then we can infer a general (shared) executive function component. Similar logic was used to identify a general attention component that explained performance across diverse tasks involving attention (Rosenberg et al., 2016; Rosenberg et al., 2019; Yoo et al., 2022).

Secondly, to further address the limitation that individual behavioral measures do not purely reflect the intended cognitive constructs, we developed measures of general and specific EF. General EF, corresponding to unity, can be estimated by extracting or regressing out common variance of the three EF components (Friedman & Miyake, 2017; Miyake et al., 2000). Specific EF, corresponding to the diversity of EF, can be estimated by the task-specific residual variance. CPM analysis on these new measures confirms the existence of a general and specific EF components and their connectome basis. Also, going beyond past work, our models are the first that can predict general and specific EF performance in novel participants not used in training.

## Method

### FMRI Data

The data for this project comes from the WU-Minn Human Connectome Project (HCP) (Van Essen et al., 2012) S1200 Release of February 2017. Among the various neuroimaging modalities available in the dataset, we specifically used the 3T resting-state fMRI scans and 2-back task fMRI data. Detailed scanning parameters for all fMRI sessions are available in Van Essen et al (2012).

Our initial filtering process retained subjects who had full completion of the resting-state fMRI sessions, a set of task fMRI sessions (Working Memory, Social, and Emotion tasks; The latter two were not used in this project.) as well as the full set of out-of-scanner NIH Toolbox tasks (NIH Toolbox®; n=445 removed from this step). We discarded subjects displaying significant head motion (≥ 3mm translation, ≥3° rotation, and ≥0.15 mm mean frame-to-frame displacement), or those missing head movement parameter files for any of the resting-state or task fMRI runs (n=2 removed). Furthermore, we excluded subjects with fMRI data flagged for known defects by the HCP team (n=11 removed). Consequently, our final sample comprised n=748 (female: 418) subjects. Our sample size is comparable or larger than that of previous CPM studies (e.g., Avery et al., 2020; Yoo et al., 2022) and provides a power close to 1 even with minimal effect size and significance level (effect size=0.1, 𝛼=0.0001).

For each included participant, two resting-state sessions were available, collected across two separate days. Each session consists of two runs (about 15 minutes each) with different phase encoding directions: left-to-right (LR) and right-to-left (RL). The N-back task data was collected over two sessions, each lasting approximately 5 minutes and using 2 different phase encoding directions, the same as in the resting sessions. The N-back task session included both 2-back and 0-back blocks. Importantly, we only used the fMRI timeseries associated with the 2-back trials, since the 0-back trials do not significantly engage working memory (K. M. Miller et al., 2009).

To compare the robustness of fMRI data representations, we tried both the volumetric (NIFTI) and grayordinate (CIFTI) fMRI data for our analysis. Both types of resting-state fMRI data were processed using the HCP minimal preprocessing pipeline (Glasser et al., 2013). The volumetric data was registered to 2mm MNI space, while the grayordinate data was additionally transformed to the standard CIFTI grayordinate space. Further preprocessing of the volumetric data involved customized Python code to remove 12 motion parameters, white matter and cerebral spinal fluid (CSF) signals, global signals, and linear trends in the timeseries data. For the resting-state grayordinate fMRI data, we used a version further denoised by an additional ICA-FIX procedure (Salimi-Khorshidi et al., 2014), which enhances the signal-to-noise ratio by isolating and removing independent components linked to motion and other artifacts. Additionally, white matter and CSF signals, global signals, and linear trends were regressed out for consistency with the volumetric preprocessing.

The 2-back volumetric fMRI data underwent the same preprocessing procedures as the resting-state data. However, for the 2-Back grayordinate data, ICA-FIX was not applied due to insufficient data to train the denoising classifier. Instead, we regressed out the 12 motion parameters as done in the volumetric case, followed by the same nuisance variable regression for white matter and CSF, global signal, and linear trend.

### Behavioral Data

The behavioral data for this project were derived from the performance measures for the 2-back task, the Dimensional Change Card Sort Test (DCCS), and the Flanker task from the HCP dataset. Every subject included in our neuroimaging sample had all 3 behavioral scores available. The 2-back task was conducted inside the scanner during the working memory session, whereas the DCCS and Flanker tasks were completed outside during a “NIH Toolbox Behavioral Tests” session. Both accuracy and response time were recorded for all 3 tasks and integrated into a normalized score for each subject. The scores for DCCS and Flanker were provided by the HCP team and preprocessed in accordance with the procedures detailed in the NIH Toolbox Scoring and Interpretation Guide found in the reference section. For the 2-back task, we processed the 2-back accuracy and response time for each individual following the same procedure using our customized Python code. The only modification we made was to lower the accuracy threshold from 4 to 2 when combining accuracy and threshold, to ensure data normality. All measurements were normalized to have a mean of 100 and a standard deviation of 15, following the standard normalization procedure outlined in the NIH Toolbox manual. Note that the out-of-scanner List Sorting scores were excluded as a measure of Updating since the task did not record response time data.

### Connectome-based Predictive Modelling

Connectome-based Predictive Modelling (CPM) (Finn et al., 2015; Shen et al., 2017) links individual differences in brain functional connectivity and behavior measures.

To construct the functional connectivity matrices, we utilized the Shen268 (Shen et al., 2013) whole-brain atlas for parcellating the volumetric data, and Schaefer300 atlas (Schaefer et al., 2018) for parcellation of the cortical part of the grayordinate data. We selected Shen268 atlas for its extensively use in previous CPM analysis on volumetric fMRI data (Avery et al., 2020; Beaty et al., 2018; Finn et al., 2015; Rosenberg et al., 2016; Yoo et al., 2018) and Schaefer300 atlas for its comparable parcellation size in the cortical surface space. Subcortical parcels in the grayordinate data were adopted from the original CIFTI labels. The average timeseries within each ROI was computed to represent the activity at that node, and pairwise Pearson correlation of all nodes was used to generate the functional connectivity matrix for each subject. Each Pearson r value underwent Fisher transformation to obtain a z value. For ease of calculation, we vectorized each individual’s connectivity matrix and concatenated them to form the connectivity matrix for the entire sample. It is important to note that each scanning session (e.g., REST1, REST2, WM) yields two connectivity matrices corresponding to the two different phase encoding runs (L-R and R-L). We later averaged these two matrices to produce a single connectivity matrix for that session. In addition, for resting-state functional connectivity data, since there are two resting-state sessions per person, we further averaged the two to obtain a single resting-state functional connectivity matrix for each individual.

### CPM Prediction of Executive Functions

We constructed CPMs for the three EF components – Inhibition, Shifting, and Updating – by using behavioral measures from the Flanker task, the Dimensional Change Card Sort (abbreviated as “Card Sort” below) task, and the 2-back task, respectively.

The CPM approach involves two principal stages: feature selection and model fitting. During feature selection, we used Pearson’s correlation to associate the connectivity edges with the behavior measure, identifying the correlation score between each edge and the selected behavior. Only edges surpassing the significance threshold (*p*<0.01, two-sided) were kept for model fitting. This process distinguishes two types of edges for selection: positively-associated edges (referred to as “positive edges” hereafter) and negatively-associated edges (“negative edges”), based on the sign of their correlation with the behavior scores. To control for variations in parcellation atlas sizes and further constrain the model to the most predictive edges, we retained only the top 100 significant positive and negative edges for subsequent analysis. Next, to fit the model, we summed together the selected positive and negative edge weights for each participant to generate an aggregate positive score and an aggregate negative score, respectively. Three linear regression models are then developed: one using the positive score only (“positive model”), one using the negative score only (“negative model”), and one combining both scores (“both model”).

To evaluate predictive performance without overfitting, we employed a 10-fold cross-validation method. Specifically, we shuffled and divided the data into 10 equal parts, training the CPM on 9 of them and testing it on the remaining one. This procedure is iterated 10 times, ensuring each fold is used for testing exactly once. The training set is utilized for feature selection and model fitting, whereas the testing set is for assessing the model’s performance. Pearson’s correlation between the predicted and actual behavioral scores is computed to gauge the model’s prediction accuracy. The average correlation across all 10 folds serves as the final measure of model fit. This process was repeated 1,000 times, with a different shuffle each time, to confirm the reliability and reproducibility of our findings. The mean model fit score across these 1,000 iterations was reported as the model’s final score. Additionally, a permutation test follows the same steps but with the subject’s behavioral scores randomly shuffled before being split into folds, ensuring a rigorous evaluation of model performance.

We assessed the within-task prediction accuracy of each CPM – evaluating the prediction for the same task on which the model was trained. This approach allows us to directly ascertain whether the individual differences in functional connectivity identified by the CPM model are robust enough to accurately predict new data.

In addition, we explored cross-prediction across every pair of tasks to evaluate the generalizability of each task-specific CPM model to predict individual differences in a different task. Cross-predictions also allow us to infer the separability of different EF components, namely what’s common and what’s specific across tasks. To this end, we tested the model on a different measure from what it was being trained on. For cross-validation, during each iteration, we maintained the exact same train-test split and applied the previously trained CPM model to predict a different task measure within the testing cohort.

### CPM Canonical Network Analysis

We also examined the anatomy of the features selected across all models to evaluate the influence of specific anatomical/functional networks on the three cognitive measures. Due to the variability in edge selection across iterations of CPM training resulting from different training-testing splits, we included an edge only if it was selected in more than half of the total iterations (over 500 out of 1000 iterations). This approach helps preserve only core edges that reflect meaningful individual differences in behavior. To confirm that the choice of threshold did not qualitatively change the results, we varied the thresholds between 40%-60%, but did not see substantial changes of the edges within or between canonical networks.

To define canonical networks on volumetric data, we used the 10-network version of the Shen 268 atlas, where each parcel is categorized into one of the medial frontal, frontoparietal, default mode, motor, visual A, visual B, visual association, salience, subcortical, cerebellum networks. For grayordinate data, we employed the 7-network version of the Schaefer 300 atlas for cortical regions. For subcortical structures, we divided them into two separate networks based on their original label given by the CIFTI file: a subcortical network and a cerebellum/brainstem network. Furthermore, since the subcortical network includes regions such as the hippocampus and thalamus, which are traditionally classified within the limbic network, we combined these regions with the limbic network as defined in the Schaefer 300 atlas and retained the designation “subcortical network” for consistency.

### CPM Computational Lesion Analysis

To further assess a canonical network’s contribution and importance to prediction performance, we performed a computational lesion analysis (also known as an ablation analysis). In this approach, we removed all connectivity patterns associated with a specific network and evaluated how this affected the predictive performance of CPM. Specifically, for each target network under investigation, we retrieved the CPM model trained using the standard procedure, removed all within and cross-connectivity edges associated with the target network, and then predicted outcomes on the hold-out testing set using the same cross-validation procedure.

### General and Specific Executive Functions Analysis

We conducted additional analyses to address the following two questions. First, is there an overarching functional connectome for general EF (Unity; Friedman & Miyake, 2017)? Second, given that the original measures may not purely reflect the underlying EF constructs (as indicated by their high correlation), can we identify the functional connectome unique to each construct (Diversity; Friedman & Miyake, 2017)?

To address these questions, we devised two types of measures: a general EF score and three component-specific scores of Inhibition, Shifting, and Updating. The general EF score was derived from the mean of the original z-scored task measures, where we hope to retain only the core information shared by different EF components. On the other hand, the component-specific scores were defined as the residuals after regressing out the other two measures from each target measure, with the aim to preserve only the variance that cannot explained by other measures. We created CPMs for the general EF score and the three component-specific scores.

## Results

### The Raw Executive Function Measures Are Correlated

To investigate the EF components of response inhibition, set switching, and working memory updates, we used behaviors from the Flanker task, the Card Sorting task, and the 2-back tasks, respectively.

Descriptive statistics revealed that every pair of measurements were correlated with each other (with *p*<0.001), as indicated in Table 1. The highest correlation was found between the Card Sort and Flanker measures. More statistics for each measurement can be found in Supplementary Table 1.

**Table 1.** Pearson Correlation between behavior measures.

|  | Flanker | Card Sort | 2-back |
| --- | --- | --- | --- |
| Flanker | -- | -- | -- |
| Card Sort | 0.4905*** | -- | -- |
| 2-back | 0.3094*** | 0.3525*** | -- |

\*\*\* $p < 0.001$ , uncorrected

### CPMs Predict Individual Differences for 3 Executive Function Tasks

Table 2 shows the results of CPMs trained and tested on Flanker, Card Sort, and 2-back task data using connectomes constructed based on 2-back task (left column) or resting-state fMRI data (right column). Most predictions were statistically significant (Permutation test *p*<0.05, corrected for FWE). The three rows of tables correspond to CPMs based on positive, negative or both types of edges. For instance, the top left entry (0.1443) in the first sub-table corresponds to the predictive performance of a positive CPM model, when trained and tested both on Flanker measures using resting-state functional connectivity.

**Table 2.** CPM prediction accuracy of raw EF measures, grayordinate.

|  |  | *Working Memory (2-back)* |  |  | *Rest* |  |  |
| --- | --- | --- | --- | --- | --- | --- | --- |
|  |  | Flanker | Card Sort | 2-back | Flanker | Card Sort | 2-back |
| Positive | Flanker | 0.23*** | 0.18*** | 0.35*** | 0.16<sup>*,a</sup> | 0.13<sup>†,b</sup> | 0.32*** |
|  | Card Sort | 0.20*** | 0.26*** | 0.35*** | 0.16<sup>*,c</sup> | 0.21*** | 0.26*** |
|  | 2-back | 0.24*** | 0.23*** | 0.42*** | 0.21*** | 0.18*** | 0.37*** |
| Negative | Flanker | 0.28*** | 0.20*** | 0.34*** | 0.20*** | 0.13<sup>†,d</sup> | 0.27*** |
|  | Card Sort | 0.25*** | 0.31*** | 0.36*** | 0.16<sup>*,e</sup> | 0.20*** | 0.25*** |
|  | 2-back | 0.28*** | 0.27*** | 0.44*** | 0.23*** | 0.18*** | 0.35*** |
| Both | Flanker | 0.29*** | 0.21*** | 0.36*** | 0.20*** | 0.14<sup>*,f</sup> | 0.31*** |
|  | Card Sort | 0.25*** | 0.32*** | 0.39*** | 0.18*** | 0.23*** | 0.29*** |
|  | 2-back | 0.29*** | 0.28*** | 0.48*** | 0.25*** | 0.20*** | 0.39*** |

Each sub-table (bounded by bold lines) represents a set of CPM prediction performance scores for a specific fMRI task state (2-back working memory vs. rest) using one type of edges (positive, negative, or both). For example, the top-left sub-table shows the CPM performance using working memory task fMRI data and only positive-associated edges. In each 3x3 sub-table, each row represents the training behavior, and each column represents the testing behavior. The maximums of each row and column in the sub-tables are colored in blue and yellow, respectively, with the overlap colored in green. \*\*\* $p < 0.001$ , \*\* $p < 0.01$ ; \* $p < 0.05$ ; † $p < 0.1$ , all corrected for family-wise error (FWE) after permutation testing. Exact p-values: <sup>a</sup>: $p = 0.012$ ; <sup>b</sup>: $p = 0.060$ ; <sup>c</sup>: $p = 0.024$ ; <sup>d</sup>: $p = 0.072$ ; <sup>e</sup>: $p = 0.012$ ; <sup>f</sup>: $p = 0.012$ .

Within-task predictions, as represented by the diagonal of each table, were significant in all cases. Notably, 2-back performance was predicted significantly more accurately than Flanker and Card Sort scores (*p*<0.05, corrected for FWE), regardless of brain state. More interestingly, cross-task prediction performance (that is, train the CPM on one behavioral measure and test on another) was also significant in most cases (permutation test *p*<0.05, corrected for FWE). 2-back performance was most strongly cross-predicted when trained on any of the task measures (*p*<0.05, corrected for FWE), and the model trained on 2-back behavior scores also showed higher cross-prediction accuracy on both Card Sort and Flanker scores. On the other hand, cross-prediction performance for Card Sort and Flanker (both ways) was numerically lower and even non-significant when using resting-state fMRI data.

Comparing the left and right columns shows that CPM performance was significantly higher from 2-back task scans than from resting-state scans (p’s <0.001 in all cases, corrected for FWE). In other words, the 2-back task fMRI functional connectivity data enabled higher prediction 𝑟 (permutation test *p*<0.05, corrected for FWE) across all model types and behavioral combinations.

We also examined the impact of fMRI data format on CPM performance by comparing models trained and tested on grayordinate data versus volumetric data. Grayordinate-based data showed superior CPM predictive performance over traditional volumetric-based data in most cases (*p*<0.001 in all cases using 2-back fMRI data; *p*<0.001 in 18 out of 27 cases using resting-state fMRI data, both corrected for FWE). Detailed CPM performance on volumetric fMRI data is presented in Supplementary Table 4.

Therefore, due to the greater variance captured by using grayordinate-based task fMRI data, we will primarily present the results based on 2-back grayordinate fMRI data in the following sections. For those interested in the results on resting-state or volumetric data, which produced comparable patterns of results, please refer to the supplementary tables for further details.

### The FPN, DMN, and DAN Engage Across All Executive Function Tasks

We next investigated the connectome anatomy for each of the EF task measures. The resulting connectome profile for each task is depicted in Figure 1. In summary, positive predictive edges for the Flanker task were mainly within the frontoparietal network and with the dorsal attention network and cerebellum. Positive predictive edges for the Card Sort task involved mainly the frontoparietal, default mode, and salience networks. Lastly, positive predictive edges for 2-back performance were mostly within the frontoparietal network, as well as between the frontoparietal network and both the default mode and dorsal attention networks. Overall, the edges that positively predicted the three EF measures span a wide range of canonical networks and are relatively distinct from one another. The core cluster of positive edges common to all three EF CPMs, illustrated in Figure 1 (bottom right), was located within the frontoparietal, and between the frontoparietal, dorsal attention, and default mode networks.

**Figure 1.**
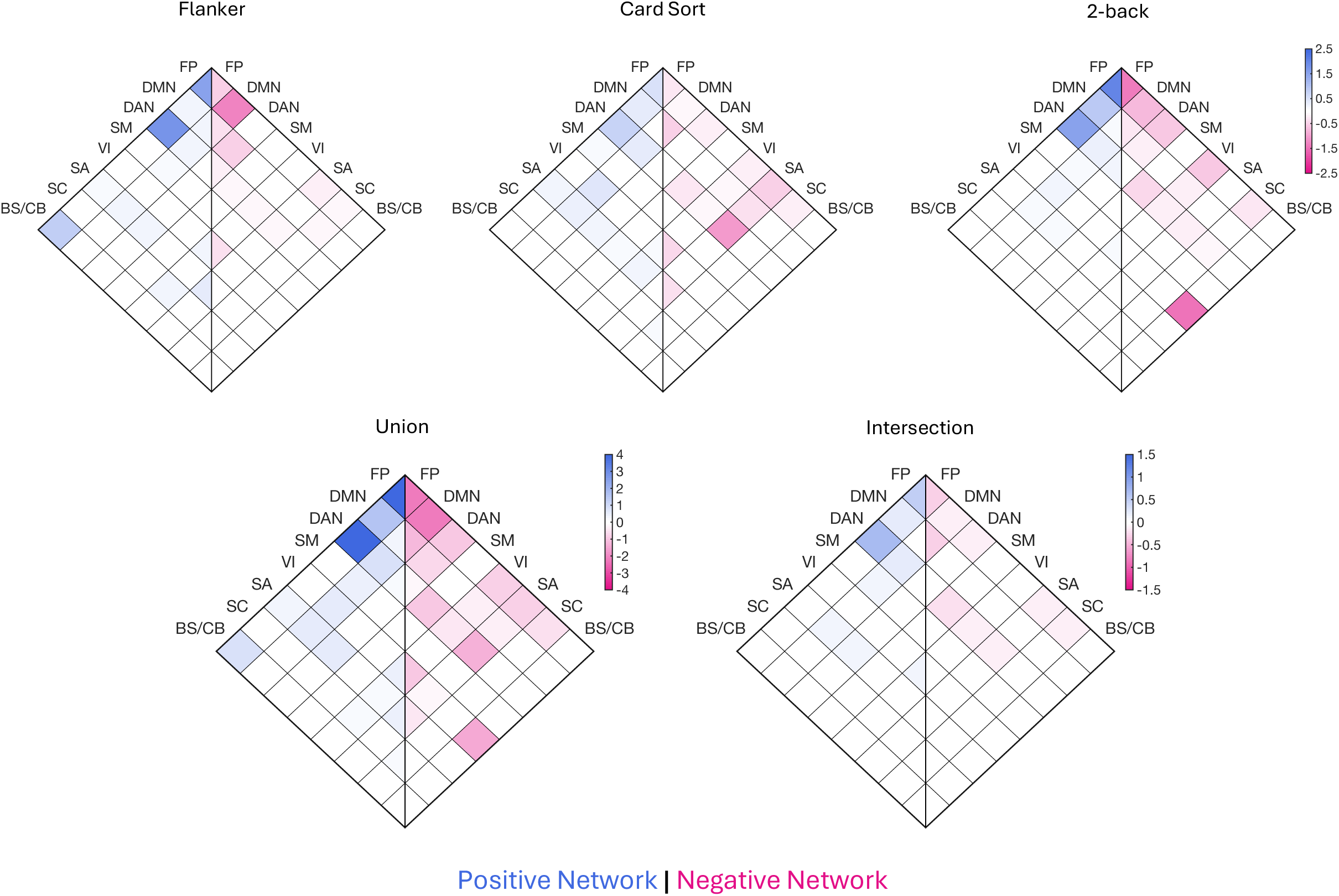
Predictive edges identified by CPM on three EF measures. The top row displays the edges identified by CPMs trained on each of the three types of EF measures. Each heatmap is divided into left and right halves, where the right half represents negative edges, and the left half represents positive edges. Each cell within the heatmap indicates the percentage of reliable edges (see the Methods section for detailed inclusion criteria) picked up by CPM between the corresponding pair of canonical networks. Higher percentages are represented by more intense pink or blue hues. The bottom row illustrates the union (sum) and intersection (minimum) of the three heatmaps above. FP, frontoparietal; DM, default mode; DA, dorsal attention; SM, somatomotor; VI, visual; SA, salience; SC, subcortical; BS/CB, brain stem/cerebellum.

On the other hand, CPM also picked up a set of negative edges that inversely relates to performance in each EF task. For the Flanker task, negative edges were found within and between the frontoparietal, default mode, and dorsal attention networks, as well as some edges within the visual network. The CPM for the Card Sort task included negative edges within the default mode, visual, and salience networks, as well as between the salience network and the dorsal attention and frontoparietal networks. The negative CPM model for the 2-back task included mostly edges associated with the frontoparietal network, along with the those between the visual network and cerebellum. The intersection plot (Figure 1, bottom right) shows that most of the negative edges common to all three CPMs were within frontoparietal and default mode networks, as well as between somatomotor and default mode networks.

### Lesioning the FPN and DMN Led To Largest Performance Drop

To further study the contribution of each network in predicting individual differences, we then performed computational lesion analysis on CPM by lesioning each network’s connectivity one at a time. As shown in Figure 2, in the positive models, lesioning frontoparietal connectivity resulted in the largest prediction performance drop in most cases, where lesioning the default mode network showed the largest prediction performance drop in the other cases. In the negative models, most impairments can be attributed to lesioning the frontoparietal network and especially the default mode network. To a lesser extent, lesioning other networks such as the salience and visual networks resulted in a significant drop in prediction performance.

**Figure 2.**
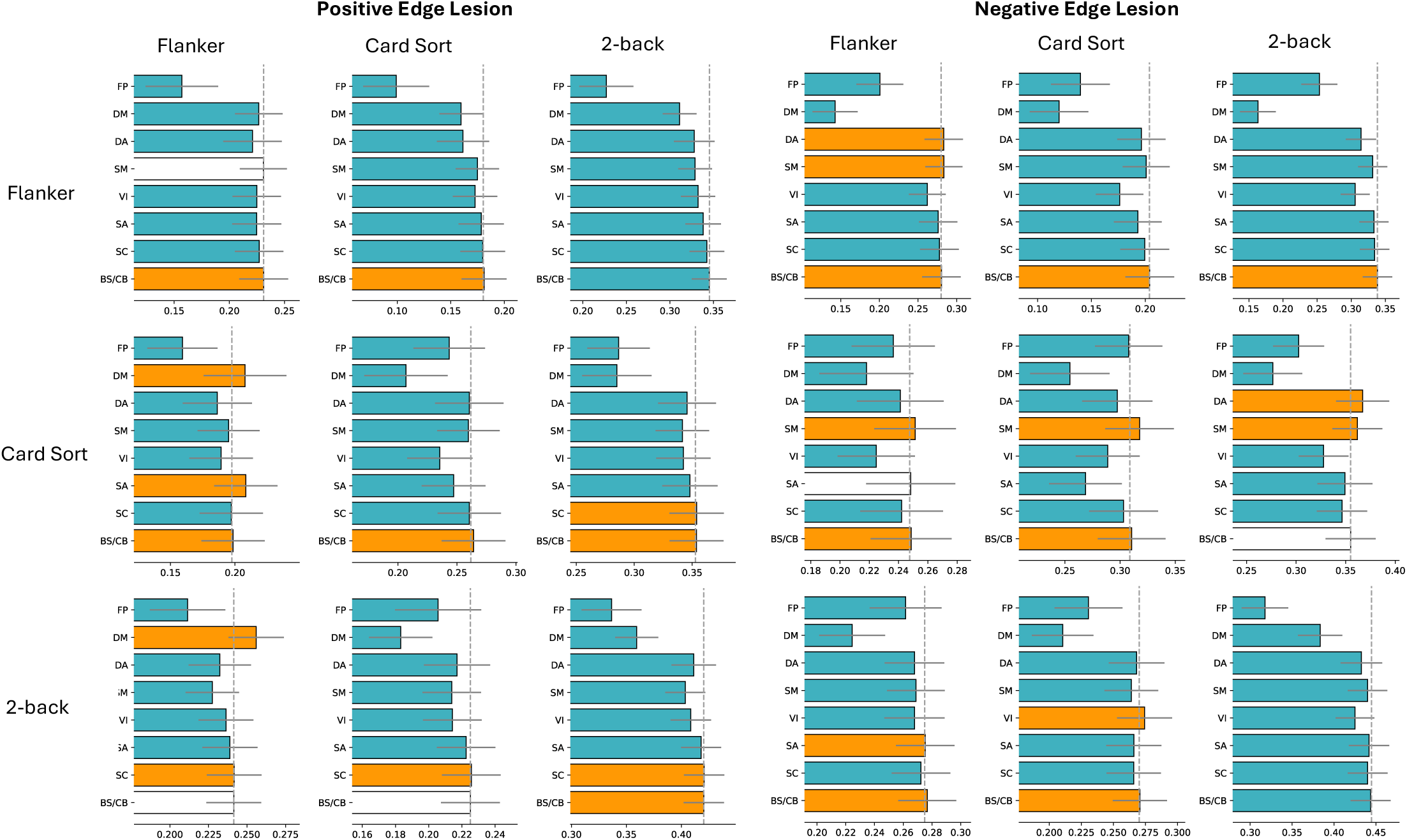
CPM lesion analysis on grayordinate, 2-back task-fMRI data. The figure is divided into two main sections: the left half displays CPM performance after lesioning positive edges of each network, while the right half shows the results after lesioning negative edges. Each section is further divided into subplots based on different train/test task combinations. In these subplots, each row corresponds to a training task, and each column represents a testing task. For example, the top-left subplot in the left half illustrates CPM prediction when trained and tested both on the Flanker scores after lesioning positive edges of each canonical network. The horizontal gray line attached to each bar denotes the 95% (± 2 standard deviation) confidence interval. The vertical gray dashed line represents the baseline CPM performance (without lesioning) for each scenario. To assess the impact of lesioning, paired t-tests were conducted comparing the lesioned performance to the regular performance. The results are depicted with colored bars: blue bars indicate significant reductions in performance (*p*<0.05, after FWE correction), orange bars represent significant increases, and white bars denote no significant difference from regular performance. All significant bars except the within-Card Sort prediction in the frontoparietal negative edge lesion condition have *p*<0.001 (FWE corrected). The within-Card Sort prediction when lesioning FP negative edges has *p* = 0.037. For non-significant conditions (white bars): Flanker-Flanker, SM, positive (*p*=5.96); 2Back-Flanker, BS/CB, positive (*p*=43.17); 2Back-Card Sort, BS/CB, positive (*p*=0.078); Card Sort-Flanker, SA, negative (*p*=0.37); Card Sort-2Back, BS/CB, negative (*p*=15.30). FP, frontoparietal; DM, default mode; DA, dorsal attention; SM, somatomotor; VI, visual; SA, salience; SC, subcortical; BS/CB, brain stem/cerebellum.

### General Executive Function is Predicted Better Than Specific Executive Function Components

To study the shared connectome underlying general EF, as well as the idiosyncratic connectomes for specific EF components, we repeated the CPM procedure on four newly devised measures: one general EF score and three component-specific scores. The general EF score was derived from the mean of the original z-scored task measures, and the component-specific scores were defined as the residuals after regressing out the other two measures from each target measure. More statistics for each measurement can be found in Supplementary Tables 2 and 3.

As illustrated in Table 3, when tested on the same measure as the CPM was trained on, the general EF measure was predicted better than the component-specific measures using 2-back fMRI data. Also, after regressing out other non-target measures, the cross-task prediction performance between different component-specific measures was dampened (*p*<0.05 in all 18 cases, corrected for FWE).

**Table 3.**
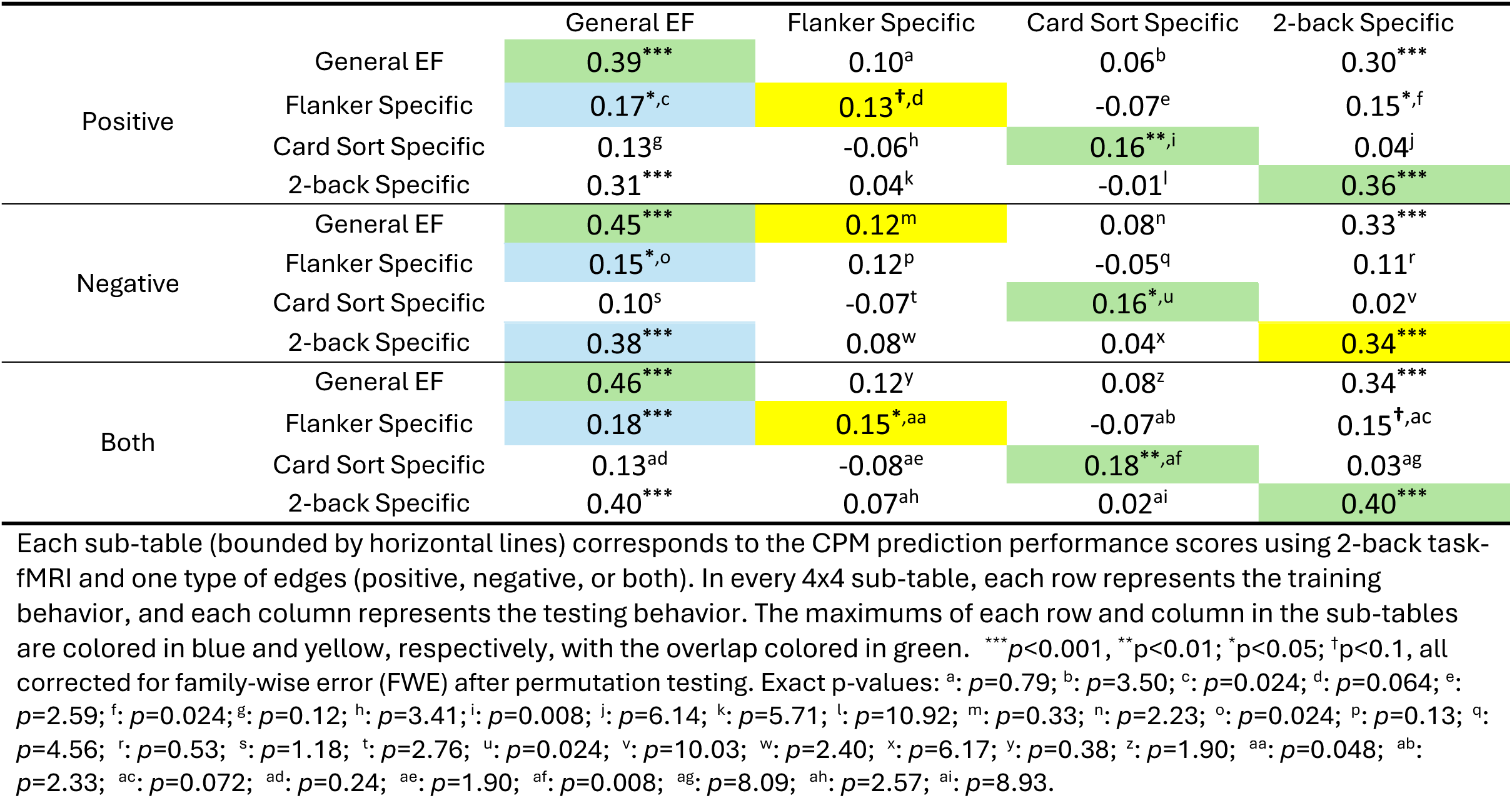
CPM prediction accuracy of general and specific EF measures, grayordinate.

Aligned with our earlier analysis, using 2-back fMRI data enhanced predictive performance over the resting-state data. Therefore, the following sections will emphasize results obtained from task-based fMRI data over resting-state fMRI data. More detailed results for the other conditions are provided in the supplementary tables and figures.

### The Component-specific Measures Bear Little Network Connectivity Overlap

Examining the canonical functional networks for each component-specific positive CPM model reveals little overlap Figure 2 (bottom right), suggesting that the connectome for each EF component became more unique when shared variance was removed.

While the connectome for each component looks roughly similar to the previous ones, one can spot some differences relative to the models without shared variance removed (Figure 1). For positive networks, the Flanker-specific component model now consists of fewer frontoparietal-related edges, both in terms of within frontoparietal edges and between frontoparietal and other networks such as dorsal attention network and brainstem/cerebellum. Within-salience network connectivity was more relevant to predict individual differences in performance (all p<0.001, FWE corrected). Similar patterns were observed for the Card-Sort-specific component model, where the frontoparietal network showed fewer connections within itself and with the dorsal attention network, while the salience network’s connections were increased (p<0.001, FWE corrected). Lastly, the 2-back-specific component model showed fewer positive edges between dorsal attention and frontoparietal networks, but more edges between within dorsal attention and between dorsal attention and default mode networks (p<0.001, FWE corrected).

Negative CPM models in Figure 3 also showed a decrease in overlapping edges relative to the models without shared variance removed (Figure 1). Again, subtle changes can be observed when closely examining each component CPM. For the Flanker-specific component model, the reduction in within-frontoparietal edges was accompanied by an increase in the interconnection between the salience and default mode networks (p<0.001, FWE corrected). The Card-Sort-specific component model showed more within-frontoparietal and brainstem/cerebellum-frontoparietal edges (p<0.001, FWE corrected). Additionally, the 2-back-specific component model showed an increase in visual-frontoparietal edges (p<0.001, FWE corrected).

**Figure 3.**
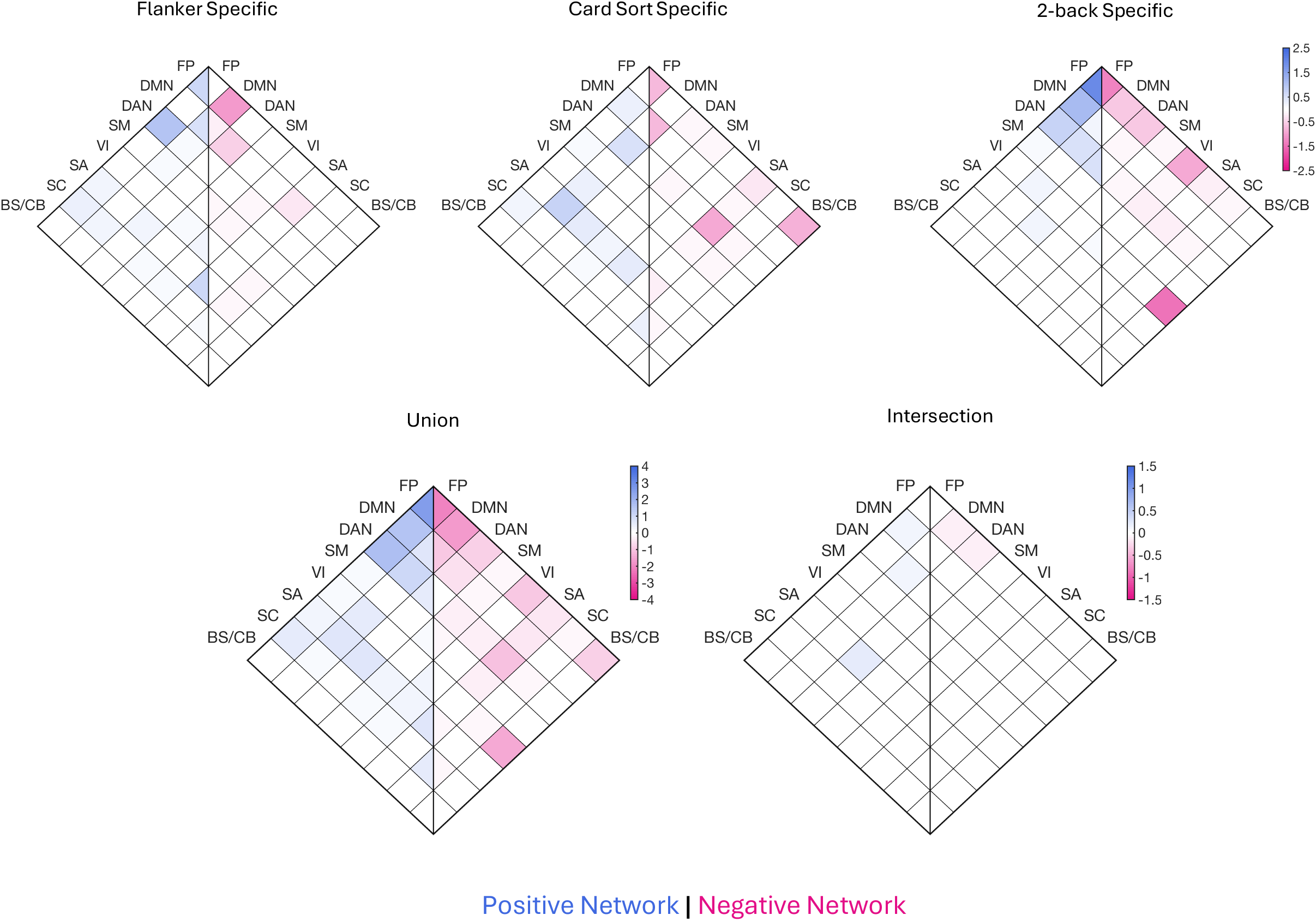
Predictive edges identified by CPM on the three component-specific measures. The top row displays the edges identified by CPMs trained on each of the three component-specific measures. The bottom row illustrates the union (sum) and intersection (minimum) of the three heatmaps above. FP, frontoparietal; DM, default mode; DA, dorsal attention; SM, somatomotor; VI, visual; SA, salience; SC, subcortical; BS/CB, brain stem/cerebellum.

### General Executive Function Involves the Interplay of the FPN, DMN, and DAN

Lastly, we examined the connectome for the general EF measure, defined as the shared variance across the Flanker, Card Sort, and 2-back tasks. As shown in Figure 4, the positive edges were densely located around the frontoparietal networks, including within-network connections and its interconnections with the dorsal attention and default mode networks. Other edges, though less densely populated, mostly bridged the default mode network with other networks. In contrast, the negative edges were most numerous within the frontoparietal network, between the frontoparietal and default mode networks, and between the cerebellum and visual networks. Additionally, the dorsal attention and visual networks comprised a substantial number of inter-network connections that inversely contributed to individual behavioral differences. These findings closely align with the intersection plot in Figure 1, reinforcing its validity as the connectome for general EF.

**Figure 4.**
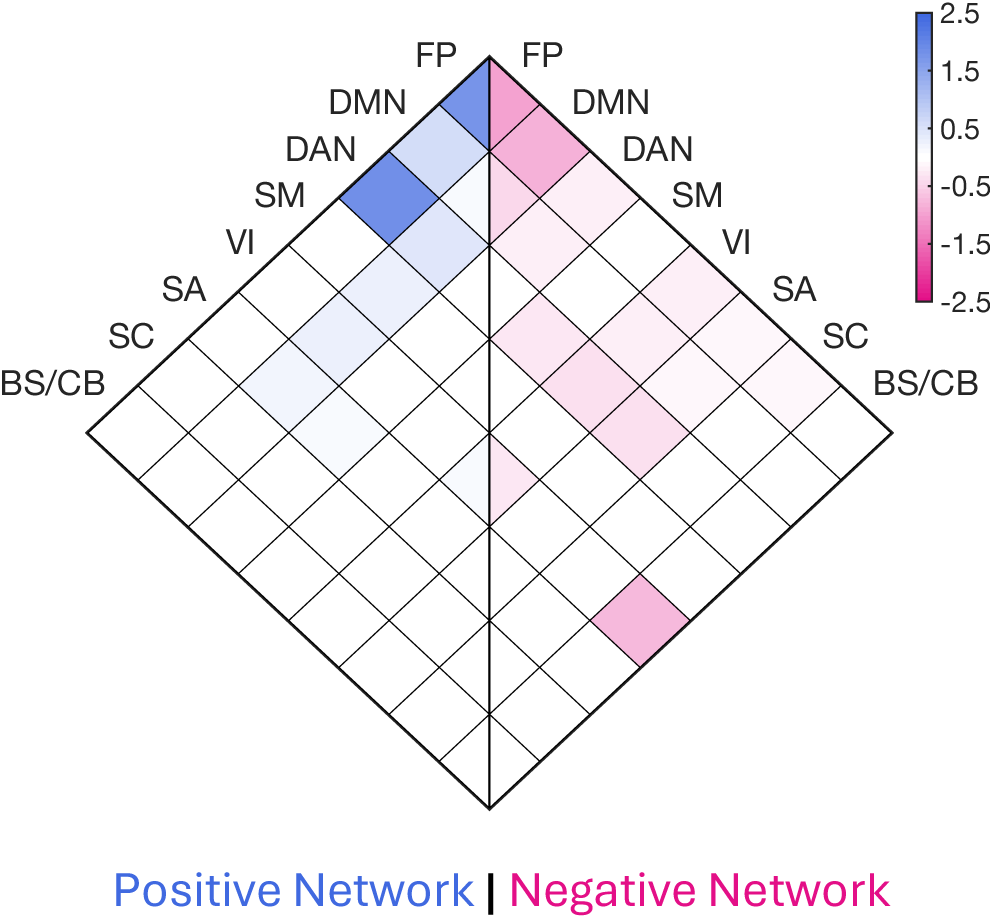
Predictive edges identified by CPM on the general EF measure. FP, frontoparietal; DM, default mode; DA, dorsal attention; SM, somatomotor; VI, visual; SA, salience; SC, subcortical; BS/CB, brain stem/cerebellum.

## Discussion

In this study, we identified both shared and specific executive function components and their brain connectomes. Utilizing Connectome-based Predictive Modeling (CPM), we revealed the distributed functional connectivity patterns that robustly predicts individual differences and neural connectome profiles underlying the Flanker task, the Dimensional Change Card Sort task, and the 2-back task. These tasks were chosen from the HCP dataset to reflect the EF components of Inhibition, Shifting, and Updating (Friedman & Miyake, 2017), although we do not assume that each task purely measures its corresponding EF component. Accordingly, we identified both shared and specific EF connectomes through the use of cross-task prediction analysis and derived EF measures.

### Cross-prediction Performance Suggests that the 2-back Task and Updating are more Central to General Executive Function

CPM cross-prediction patterns on the three original task measures revealed that the task CPM trained on 2-back behavioral scores generalized the best to predict the performance in the other two tasks. The high cross-prediction performance suggests that the connectome basis of the 2-back task also tracks variations in Flanker and Card Sort tasks. On the other hand, the Flanker and Card Sort tasks appear to be more independent from the connectome standpoint, evident by their lower cross-prediction accuracy. It is worth noting that the higher cross-prediction accuracy of the 2-back model is not merely due to using the same fMRI and behavioral task, since the cross-prediction gain of CPMs based on 2-back scores persists with resting-state fMRI data.

These results are consistent with a central role of Updating in EF -- a notion supported by various prior studies. In Lemire-Rodger et al. (2019), multivariate analysis on fMRI data alluded to Updating as a common factor supporting the other EF processes. In another meta-analysis, Rodríguez-Nieto et al (2022). reported that the Updating network highly overlaps with the Shifting and Inhibition networks, while the latter two exhibited minimal overlap. At the behavioral level, Updating stood out as the sole process among the three EF mechanisms showing a substantial correlation with general intelligence (Friedman et al., 2006), suggesting it is domain-general. Updating, commonly operationalized as working memory, involves controlling attention to resist interference, a key component that scaffolds many higher-order executive processes (Engle, 2002; Engle et al., 1999; Kane & Engle, 2003; Unsworth et al., 2004).

Note that our finding is less consistent with some previous studies that posited Shifting (Dajani & Uddin, 2015) and Inhibition (Miyake & Friedman, 2012) as more central components for general EF. While we speculate that this difference may arise from the disparate level of focus (behavioral vs. neural connectome), further investigation is warranted to directly compare the results.

On the other hand, the Flanker task CPM (Inhibition) and the Card Sort task CPM (Shifting) exhibited lower generalizability when applied to predict other tasks, suggesting they reflect distinct components of EF. This proposition aligns with several previous studies (Lemire-Rodger et al., 2019; Miyake et al., 2000; Rodríguez-Nieto et al., 2022). Upon closer examination of our results using task fMRI data, we found that models trained on the Card Sort task and tested on the Flanker task performed better than those trained and tested in the opposite direction. This observation may be related to the idea that task switching is facilitated by inhibition, as smoothly transitioning to a new task set requires effectively suppressing the previous set of rules. (Davidson et al., 2006; Diamond, 2013; Koch et al., 2010).

### Derived Measures Reveal General and Specific Executive Function Connectomes

The high prediction accuracy of our CPM of the general EF measure, derived from the mean of the three tasks, as well as its significant cross-task prediction performance, indicate the presence of a general EF factor. The significant within-task prediction accuracy of the 2-back-specific component model, along with its high cross-predictive performance on the general EF factor, suggest a stronger role for the EF functions in the 2-back task, putatively Updating. Conversely, the cross-prediction accuracy between Flanker-specific and Card Sort-specific component measures were numerically lower, suggesting that they became more distinctive after removing the general aspect of EF. Overall, the identification of a general EF factor aligns with Miyake’s theory (Friedman & Miyake, 2017; Miyake et al., 2000) that posited a construct that unifies different types of executive processes.

Examining the functional networks supporting general EF, we observed that many positive edges (i.e., edges that positively correlate with performance) reside in and between the frontoparietal (FPN), dorsal attention (DAN), default mode (DMN), and salience networks (SN). All of these canonical functional networks have previously been implicated in various aspects of EF (Friedman & Robbins, 2022b; Menon & D’Esposito, 2022). The FPN is associated with the initialization and adjustment of control, executive task performance and interactions between attention and other cognitive processes (Dosenbach et al., 2008; Marek & Dosenbach, 2018; Seeley et al., 2007). The DAN directs top-down attention and assists successful spatial attention (Corbetta & Shulman, 2002, 2011; He et al., 2007). The SN plays a role in perceiving event saliency, monitoring conflicts, and initiating access to working memory and attention (Carter & van Veen, 2007; Menon & Uddin, 2010). Finally, although the DMN tends to quench its activity during tasks, it has a putative role in switching between internal and external attention modes (Leech et al., 2011). Additionally, we also found a number of positive edges in the visual network (VN). Although this may not be directly related to EF, the strength of connectivity within the VN is consistent with the fact that all three tasks involved visual perception (Baldassarre et al., 2012).

While each of these networks serve their distinct roles in EF, they communicate with each other to subserve more complicated EF processes, as reported by various prior studies. For instance, researchers have found that the FPN shows differential connectivity patterns with the DMN and the DAN. The former strengthens during cue-independent introspective tasks, while the latter relates more to perceptual attention (Dixon et al., 2018). Another study revealed that the connectivity between the DMN and other task-related networks (e.g., the SN, FPN) were strengthened during a battery of tasks and is correlated with task performances (Elton & Gao, 2015). Aligned with these results, our general EF CPM picked up these connectivity features, and thus further consolidate the notion that these functional networks may act as hubs for general EF.

Interestingly, some connectivity features that negatively correlated with general EF performance were from the same set of canonical networks. For instance, a high proportion of edges within the FPN and between the FPN and DMN appeared in both the positive and negative models. One possible interpretation is that the canonical networks such as FPN encompass a large number of brain regions that may be heterogeneous in their contributions to EF (cf. Dixon et al., 2018). As a result, the connectivity strength of different subregions of a canonical network may correlate differently with performance measures.

While many of the identified edges have established roles in prior studies, CPM also detected numerous edges that are less reported. For instance, our results revealed an inverse relationship between general EF performance and the cerebellum’s co-activation with the VN. However, we did not find reports in the literature about the cerebellum’s negative correlation with cognitive performance. Therefore, confirming or disproving this relationship warrants deliberate investigation in the future. Overall, we believe that CPM can play a valuable role in expanding our current knowledge base by generating new hypotheses for future testing.

On the other hand, the connectome profiles for each component-specific measure were more distinct from one another. For example, in the positive models, the Flanker task was characterized by a high density of edges within the FPN, DMN, and SN. In contrast, the Card Sort task showed more edges connecting the DMN with the SN and DAN. The 2-back task’s edges were primarily concentrated within and between the FPN, DMN, and DAN. These network motifs might hint at unique signatures for each specific EF component.

### Grayordinate Data Representation Enhances Prediction Accuracy

Throughout our analysis, we also studied the impact of fMRI data representation and brain state on CPM performance. We found that using grayordinate (CIFTI) data provides significant boost in CPM predictive accuracy over traditionally used volumetric (NIFTI) data. This advantage can be attributed to the inherent benefit of using grayordinate data, which registers the cortical areas into a flat surface, while maintaining 3D structures of subcortical areas. By doing so, it provides a more compact representation with higher inter-subject spatial correspondence (Glasser et al., 2013), better signal-to-noise ratio (Smith et al., 2013), and reduced signal contamination that inflates functional connectivity (Brodoehl et al., 2020). Admittedly, this is not an exhaustive comparison between the two data representations, but it suggests that a well-chosen fMRI data format can enhance CPM analysis performance. More dedicated comparisons in the future may provide deeper insights into the best practices for CPM analysis.

### Task-based Connectome Supports Better Predictions Over Resting-state

Comparing between resting-state data with 2-back task-based fMRI data, we noticed a significant improvement associated with the use of 2-back data across both volumetric and gradyordinate data representation formats. This enhancement is consistent with previous findings that task-based fMRI data generally aids in more accurate prediction of traits and behaviors, even if the task data differs from the behavior being predicted (Chen et al., 2022; Elliott et al., 2019; Finn & Bandettini, 2021; Greene et al., 2018; Jiang et al., 2020; Yoo et al., 2018; Yoo, Rosenberg, Kwon, Scheinost, et al., 2022). In combination with other reported benefits of task fMRI data, such as its greater information gain for inferring hidden parameters (Tuominen et al., 2023), less head motion during acquisition (Huijbers et al., 2017), and better test-retest reliability for network identification (Kristo et al., 2014; Rosazza et al., 2014; Wang et al., 2017), when possible, we support the use of task-based fMRI data over resting-state data (or a hybrid use of resting and task, see Finn, 2021 for further perspectives) to achieve better predictive accuracy.

### Future Directions

While our study provides a fresh perspective on the connectome profiles of various EF processes, we also consider several limitations of our current approach as well as potential future directions. First off, the so-called task impurity problem (Burgess, 1997; Phillips, 1997) poses a challenge to accurately measuring the psychological processes targeted by EF tasks. Because each EF task typically involves a combination of cognitive, perceptual, and motor processes, one cannot assert that the variance captured by the model reflects only the intended process. Given the inherent difficulty in creating a “pure” EF task, our project takes the valuable approach of distinguishing between the common and specific components that each EF score measures. Future work could build on these analyses by extending it to other cognitive and non-cognitive measures, allowing us to evaluate the extent to which each task relies on general versus specific EF components, as well as the generalizability of our EF connectome.

Secondly, due to the constrained nature of public neuroimaging datasets, we were only able to build our CPMs on the 2-back and resting-state data. While the 2-back data already demonstrates good predictive performance across all three behavior measures, it would be valuable to replicate these analyses using other types of EF-related task fMRI data. A similar concern is that the connectome profile captured using 2-back fMRI data differs from that obtained using resting-state data. Interpreting this discrepancy is essential to developing a connectome profile of EF that is agnostic to brain state.

Third, it will be informative to examine the relation between executive function and attention. As an exploratory analysis, we compared general EF here with a general attention model developed using a different set of attention tasks in a separate dataset. To predict general EF performance here, we applied an externally validated connectome model of general attention from Yoo et al. (2002) that can predict performance in gradual continuous performance tasks, multiple-object tracking, and visual short-term memory tasks. The cross-prediction accuracy on general EF was significant but numerically much lower than when predicted using general EF CPM (Supplementary Table 9a), suggesting that general EF is distinct from general attention. When applied to the three original EF measures, the general attention CPM showed numerically lower prediction accuracy than the general EF CPM (Supplementary Table 9b), reinforcing the idea that attention does not fully account for EF. Note that for consistency with the general attention CPM study, we utilized resting-state volumetric fMRI data for this analysis. Again, this comparison should be considered exploratory as the general EF model and the general attention model were trained on different data sets. Future work should test more direct comparisons.

Finally, while this study specifically examines the EF components in adults, it will be useful to explore the connectome basis of EF from a developmental perspective. Extensive studies have shown that EF and prefrontal cortex undergo prolonged developmental trajectories before stabilizing in adulthood (Anderson, 2002; Davidson et al., 2006; Diamond, 2002; Kolb et al., 2012; Luna, 2009), and the relevant functional networks exhibit significant changes to facilitate enhanced EF performance (Keller et al., 2023; Nomi et al., 2017). From the whole-brain connectivity standpoint, there are indeed shared and unique connectome features that stably predict children’s cognitive performance (Chen et al., 2022). Thus, it would be beneficial to generalize our CPM analysis pipeline to developmental cohorts, leveraging datasets like the ABCD Study® to investigate the neurodevelopmental trajectories of each facet of EF. A thorough understanding of how the connectome of EF components evolve across different ages could illuminate the origins of EF and provide insights into the neural signatures of atypical EF development.

## Conclusion

In this study, we performed connectome-based predictive modeling analysis on large-scale task fMRI data to reveal both shared and unique aspects of executive functions from a whole-brain connectivity perspective. Our analyses suggest the centrality of the Updating component in executive function and highlight the roles of the frontoparietal, default-mode, and dorsal attention networks in supporting general executive functioning. We further demonstrated the predictive advantages of using task-based grayordinate fMRI data over resting-state or volumetric data. Future work could build on this pipeline to generalize findings across additional brain states and behaviors that fall under the umbrella of executive control.

## Acknowledgments

Data were provided [in part] by the Human Connectome Project, WU-Minn Consortium (Principal Investigators: David Van Essen and Kamil Ugurbil; 1U54MH091657) funded by the 16 NIH Institutes and Centers that support the NIH Blueprint for Neuroscience Research; and by the McDonnell Center for Systems Neuroscience at Washington University. General attention connectome-based predictive model funded by National Institutes of Health, Grant Number 5R01MH108591 to M. M. C. Computing resources and S.Q. funded by Yale University.

