## Supplementary_Tables_Figures for "Connectome-based Predictive Models of General and Specific Executive Functions"

**Supplementary Table 1.** Statistics of raw behavior measures

| | Flanker | Card Sort | 2-Back |
|---|---|---|---|
| count | 748 | 748 | 748 |
| mean | 100 | 100 | 100 |
| std | 15.0100 | 15.01 | 15.01 |
| min | 58.9189 | 49.69 | -9.32 |
| 25% | 90.1104 | 89.51 | 90.63 |
| 50% | 99.3473 | 99.78 | 101.09 |
| 75% | 109.6509 | 109.56 | 109.76 |
| max | 144.86 | 140.95 | 138.20 |

**Supplementary Table 2.** Pearson Correlation between the 4 new measures

| | General EF | Flanker Specific | Card Sort Specific | 2-back Specific |
|---|---|---|---|---|
| General EF | -- | -- | -- | -- |
| Flanker Specific | 0.3730*** | -- | -- | -- |
| Card Sort Specific | 0.3671*** | -0.4287*** | -- | -- |
| 2-back Specific | 0.4006*** | -0.1674*** | -0.2422*** | -- |

***: $p<0.001$, uncorrected

**Supplementary Table 3.** Statistics of the new behavior measures

| | General EF | Flanker-specific | Card Sort-specific | 2-Back-specific |
|---|---|---|---|---|
| count | 748 | 748 | 748 | 748 |
| mean | 100 | 100 | 100 | 100 |
| std | 15.010037 | 15.010037 | 15.010037 | 15.010037 |
| min | 50.81347 | 55.554659 | 40.13981 | -17.843425 |
| 25% | 88.847471 | 89.516514 | 90.357072 | 90.915042 |
| 50% | 100.5231 | 99.330337 | 99.688935 | 101.035226 |
| 75% | 110.625526 | 109.315296 | 109.320616 | 110.210396 |
| max | 143.037699 | 156.004711 | 152.515038 | 143.44036 |

**Supplementary Table 4.** CPM prediction accuracy of raw EF measures, volumetric

| | | *Working Memory (2-back)* | | | *Rest* | | |
|---|---|---|---|---|---|---|---|
| | | Flanker | Card Sort | 2-Back | Flanker | Card Sort | 2-Back |
| Positive | Flanker | 0.16$^{*,a}$ | 0.13$^{\dagger,b}$ | 0.32$^{***}$ | 0.15$^{***}$ | 0.08$^{g}$ | 0.15$^{*,h}$ |
| | Card Sort | 0.16$^{*,c}$ | 0.21$^{***}$ | 0.26$^{***}$ | 0.06$^{i}$ | 0.15$^{*,j}$ | 0.14$^{*,k}$ |
| | 2-Back | 0.21$^{***}$ | 0.18$^{***}$ | 0.37$^{***}$ | 0.15$^{***}$ | 0.16$^{***}$ | 0.20$^{***}$ |
| Negative | Flanker | 0.20$^{***}$ | 0.13$^{\dagger,d}$ | 0.27$^{***}$ | 0.10$^{l}$ | 0.05$^{m}$ | 0.09$^{n}$ |
| | Card Sort | 0.16$^{*,e}$ | 0.20$^{***}$ | 0.25$^{***}$ | 0.07$^{o}$ | 0.15$^{***}$ | 0.19$^{***}$ |
| | 2-Back | 0.23$^{***}$ | 0.18$^{***}$ | 0.35$^{***}$ | 0.12$^{p}$ | 0.20$^{***}$ | 0.22$^{***}$ |
| Both | Flanker | 0.20$^{***}$ | 0.14$^{*,f}$ | 0.31$^{***}$ | 0.13$^{*,q}$ | 0.07$^{r}$ | 0.13$^{\dagger,s}$ |
| | Card Sort | 0.18$^{***}$ | 0.23$^{***}$ | 0.29$^{***}$ | 0.07$^{t}$ | 0.16$^{***}$ | 0.18$^{***}$ |
| | 2-Back | 0.25$^{***}$ | 0.20$^{***}$ | 0.39$^{***}$ | 0.15$^{*,u}$ | 0.20$^{***}$ | 0.23$^{***}$ |

$^{***}p<0.001$, $^{**}p<0.01$; $^{*}p<0.05$; $^{\dagger}p<0.1$, all corrected for family-wise error (FWE) after permutation testing. Exact p-values: a: 0.012; b: 0.060; c: 0.024; d: 0.072; e: 0.012; f: 0.012; g: 0.92; h: 0.012; i: 1.76; j: 0.012; k: 0.036; l: 0.22; m: 2.08; n: 0.52; o: 1.48; p: 0.13; q: 0.04; r: 1.20; s: 0.072; t: 1.16; u: 0.024.

**Supplementary Table 5.** t-test statistics for grayordinate vs. volumetric, rest

| | | Flanker | Card Sort | 2-Back |
|---|---|---|---|---|
| Positive | Flanker | -7.92$^{***}$ | 83.55$^{***}$ | 62.06$^{***}$ |
| | Card Sort | 91.19$^{***}$ | -9.13$^{***}$ | 128.05$^{***}$ |
| | 2-Back | 6.78$^{***}$ | 49.22$^{***}$ | 38.18$^{***}$ |
| Negative | Flanker | 40.95$^{***}$ | 75.49$^{***}$ | 113.57$^{***}$ |
| | Card Sort | 25.30$^{***}$ | -26.16$^{***}$ | -25.48$^{***}$ |
| | 2-Back | -6.60$^{***}$ | -14.74$^{***}$ | -33.67$^{***}$ |
| Both | Flanker | 31.19$^{***}$ | 97.35$^{***}$ | 111.44$^{***}$ |
| | Card Sort | 77.78$^{***}$ | -38.03$^{***}$ | 66.53$^{***}$ |
| | 2-Back | 14.25$^{***}$ | 6.73$^{***}$ | -1.54$^{***}$ |

$^{***}p<0.001$. a: p=0.37 (FWE corrected)

**Supplementary Table 6.** CPM prediction accuracy of general and specific EF measures, grayordinate, rest

| | | General EF | Flanker Specific | Card Sort Specific | 2-Back Specific |
|---|---|---|---|---|---|
| Positive | General EF | 0.24*** | 0.04 | 0.08 | 0.16*** |
| | Flanker Specific | 0.05 | 0.07 | -0.04 | 0.02 |
| | Card Sort Specific | 0.08 | -0.05 | 0.06 | 0.08 |
| | 2-Back Specific | 0.23*** | 0.05 | 0.08 | 0.16*** |
| Negative | General EF | 0.21*** | 0.01 | 0.09 | 0.14*** |
| | Flanker Specific | 0.02 | 0.11 | -0.07 | -0.02 |
| | Card Sort Specific | 0.06 | -0.04 | 0.06 | 0.05 |
| | 2-Back Specific | 0.20*** | 0.01 | 0.12* | 0.10 |
| Both | General EF | 0.24*** | 0.04 | 0.08 | 0.16*** |
| | Flanker Specific | 0.04 | 0.09 | -0.05 | 0.01 |
| | Card Sort Specific | 0.08 | -0.05 | 0.07 | 0.07 |
| | 2-Back Specific | 0.24*** | 0.03 | 0.10 | 0.15*** |

Each sub-table (bounded by horizontal lines) corresponds to the CPM prediction performance scores using resting-state fMRI and one type of edges (positive, negative, or both). In every 4x4 sub-table, each row represents the training behavior, and each column represents the testing behavior. The maximums of each row and column in the sub-tables are colored in blue and yellow, respectively, with the overlap colored in green. ***$p$<0.001, **p<0.01; *p<0.05; †p<0.1, all corrected for family-wise error (FWE) after permutation testing.

**Supplementary Table 7.** CPM prediction accuracy of general and specific EF measures, volumetric, 2-back

| | | General EF | Flanker Specific | Card Sort Specific | 2-Back Specific |
|---|---|---|---|---|---|
| Positive | General EF | 0.32*** | 0.08 | 0.02 | 0.29*** |
| | Flanker Specific | 0.09 | 0.06 | -0.07 | 0.12 |
| | Card Sort Specific | 0.06 | -0.04 | 0.11 | -0.01 |
| | 2-Back Specific | 0.32*** | 0.10 | 0.00 | 0.30*** |
| Negative | General EF | 0.32*** | 0.09 | 0.03 | 0.26*** |
| | Flanker Specific | 0.14† | 0.10 | -0.05 | 0.13 |
| | Card Sort Specific | 0.03 | -0.04 | 0.07 | -0.01 |
| | 2-Back Specific | 0.29*** | 0.10 | -0.01 | 0.26*** |
| Both | General EF | 0.35*** | 0.09 | 0.03 | 0.29*** |
| | Flanker Specific | 0.13 | 0.09 | -0.07 | 0.14† |
| | Card Sort Specific | 0.05 | -0.04 | 0.10 | -0.01 |
| | 2-Back Specific | 0.33*** | 0.11 | -0.01 | 0.30*** |

Each sub-table (bounded by horizontal lines) corresponds to the CPM prediction performance scores using 2-back task-fMRI and one type of edges (positive, negative, or both). In every 4x4 sub-table, each row represents the training behavior, and each column represents the testing behavior. The maximums of each row and column in the sub-tables are colored in blue and yellow, respectively, with the overlap colored in green. ***$p$<0.001, **p<0.01; *p<0.05; †p<0.1, all corrected for family-wise error (FWE) after permutation testing.

**Supplementary Table 8.** CPM prediction accuracy of general and specific EF measures, volumetric, rest

| | | General EF | Flanker Specific | Card Sort Specific | 2-Back Specific |
|---|---|---|---|---|---|
| Positive | General EF | 0.23*** | 0.07 | 0.03 | 0.16*** |
| | Flanker Specific | 0.09 | 0.10 | -0.05 | 0.06 |
| | Card Sort Specific | 0.04 | -0.07 | 0.12† | 0.00 |
| | 2-Back Specific | 0.17* | 0.04 | 0.05 | 0.11 |
| Negative | General EF | 0.22*** | -0.01 | 0.10 | 0.18*** |
| | Flanker Specific | 0.01 | 0.08 | -0.07 | 0.00 |
| | Card Sort Specific | 0.06 | -0.06 | 0.08 | 0.04 |
| | 2-Back Specific | 0.17*** | -0.01 | 0.09 | 0.12† |
| Both | General EF | 0.25*** | 0.04 | 0.07 | 0.19*** |
| | Flanker Specific | 0.07 | 0.10 | -0.06 | 0.04 |
| | Card Sort Specific | 0.06 | -0.06 | 0.09 | 0.04 |
| | 2-Back Specific | 0.18*** | 0.01 | 0.08 | 0.13* |

Each sub-table (bounded by horizontal lines) corresponds to the CPM prediction performance scores using resting-state fMRI and one type of edges (positive, negative, or both). In every 4x4 sub-table, each row represents the training behavior, and each column represents the testing behavior. The maximums of each row and column in the sub-tables are colored in blue and yellow, respectively, with the overlap colored in green. ***$p$<0.001, **p<0.01; *p<0.05; †p<0.1, all corrected for family-wise error (FWE) after permutation testing.

**Supplementary Table 9a.** Use general attention CPM to predict the general and specific EF measures, volumetric, rest, both edges

| | General EF | Flanker Specific | Card Sort Specific | 2-back Specific |
|---|---|---|---|---|
| General Attention CPM | 0.11[*, a] | 0.06 | 0.08 | 0.05 |
| General EF CPM | 0.25[***] | 0.04 | 0.07 | 0.19[***] |

The general attention CPM cross-prediction performance is calculated by correlating the predicted behavior measure with the actual behavior measure. The general EF CPM results are from Table 9. The p-values are corrected by FWE. [***]p<0.001, [**]p<0.01, [*]p<0.05; [†]p<0.1. p-values: a: 0.031.

**Supplementary 9b.** Use general attention CPM to predict the original EF measures, volumetric, rest, both edges

| | Flanker | Card Sort | 2-back |
|---|---|---|---|
| General Attention CPM | 0.08 | 0.11[*, a] | 0.09 |
| General EF CPM | 0.15[**, b] | 0.18[***] | 0.25[***] |

The model performance is calculated by correlating the predicted behavior measure (by the general attention CPM) with the actual behavior measure. The general EF CPM results are obtained via the regular CPM train/test procedure. The p-values are corrected by FWE. [***]p<0.001, [**]p<0.01, [*]p<0.05; [†]p<0.1. p-values: a: 0.031, b: 0.006.

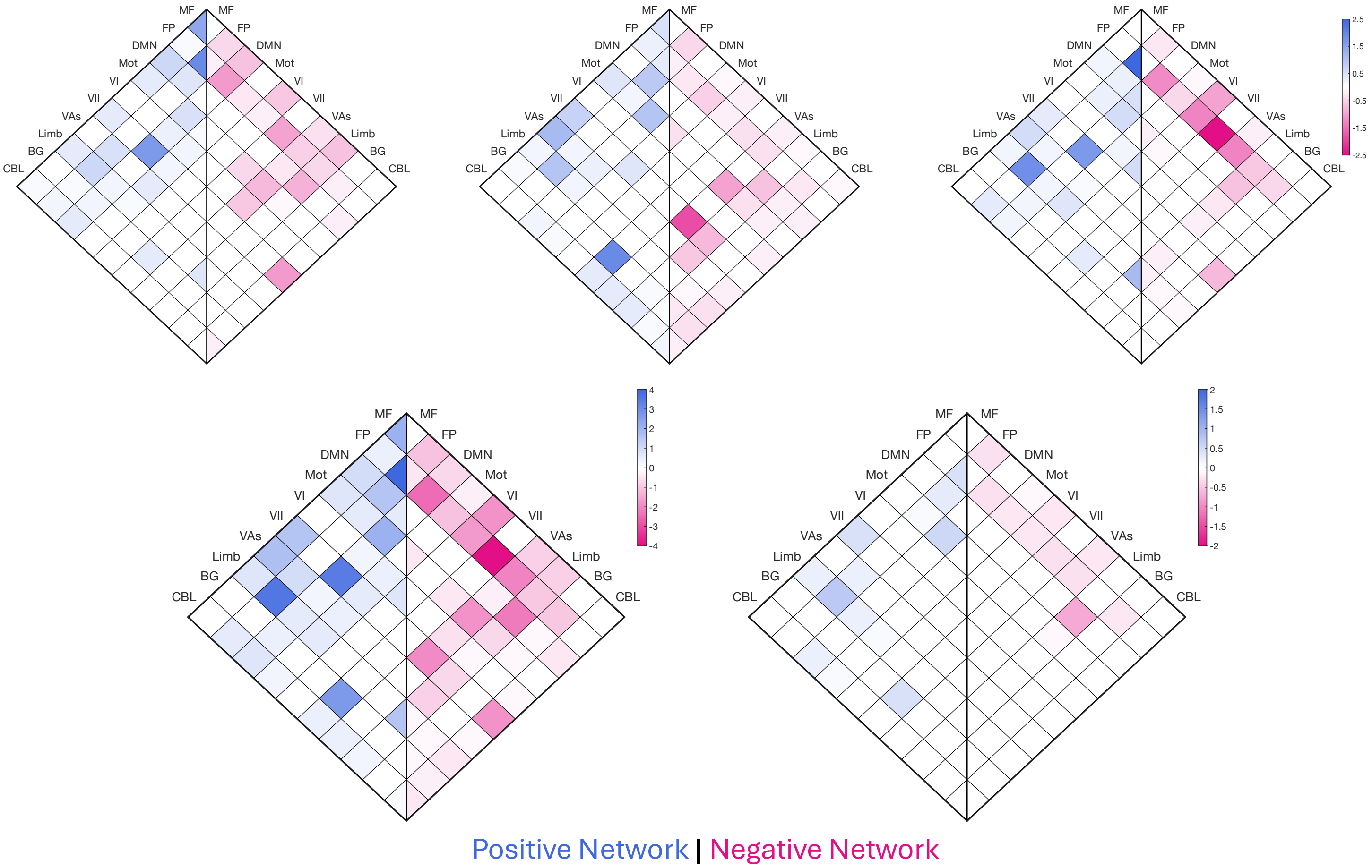


**Supplementary Figure 1.** CPM Canonical network analysis, using 2-Back, volumetric fMRI

Top 3: Flanker, Card Sort, 2-back; Bottom 2: Union, Intersection

Network Acronyms: MF, medial frontal; FP, frontoparietal; DMN, default-mode; Mot, motor; VI, visual A; VII, visual B; VAs, visual association; Limb, limbic; BG, basal ganglia; CBL, cerebellum

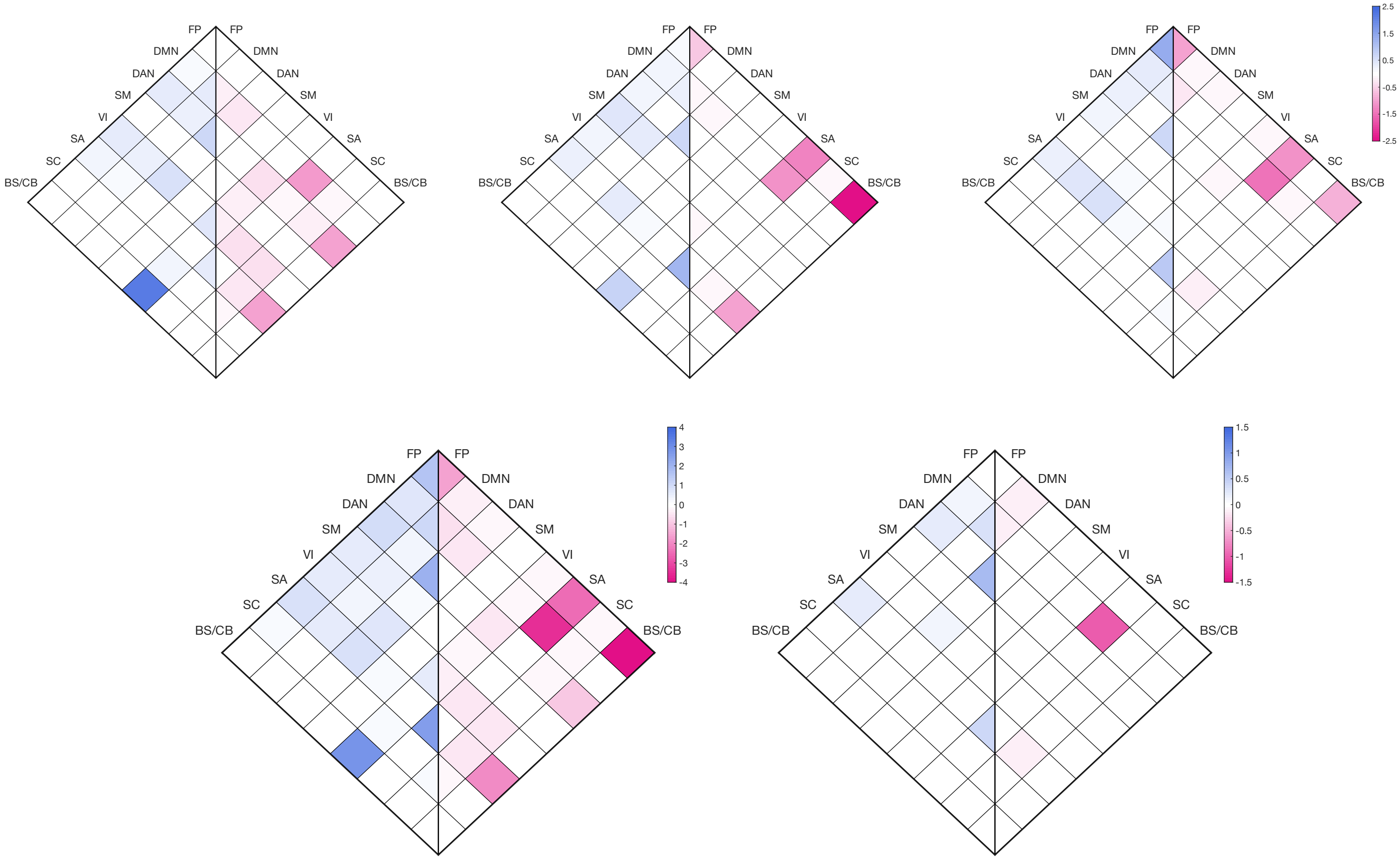


Positive Network | Negative Network

**Supplementary Figure 2.** CPM Canonical network analysis, using resting, grayordinate fMRI

Top 3: Flanker, Card Sort, 2-back; Bottom 2: Union, Intersection

Network Acronyms: FP, frontoparietal; DM, default mode; DA, dorsal attention; SM, somatomotor; VI, visual; SA, salience; SC, subcortical; BS/CB, brain stem/cerebellum.

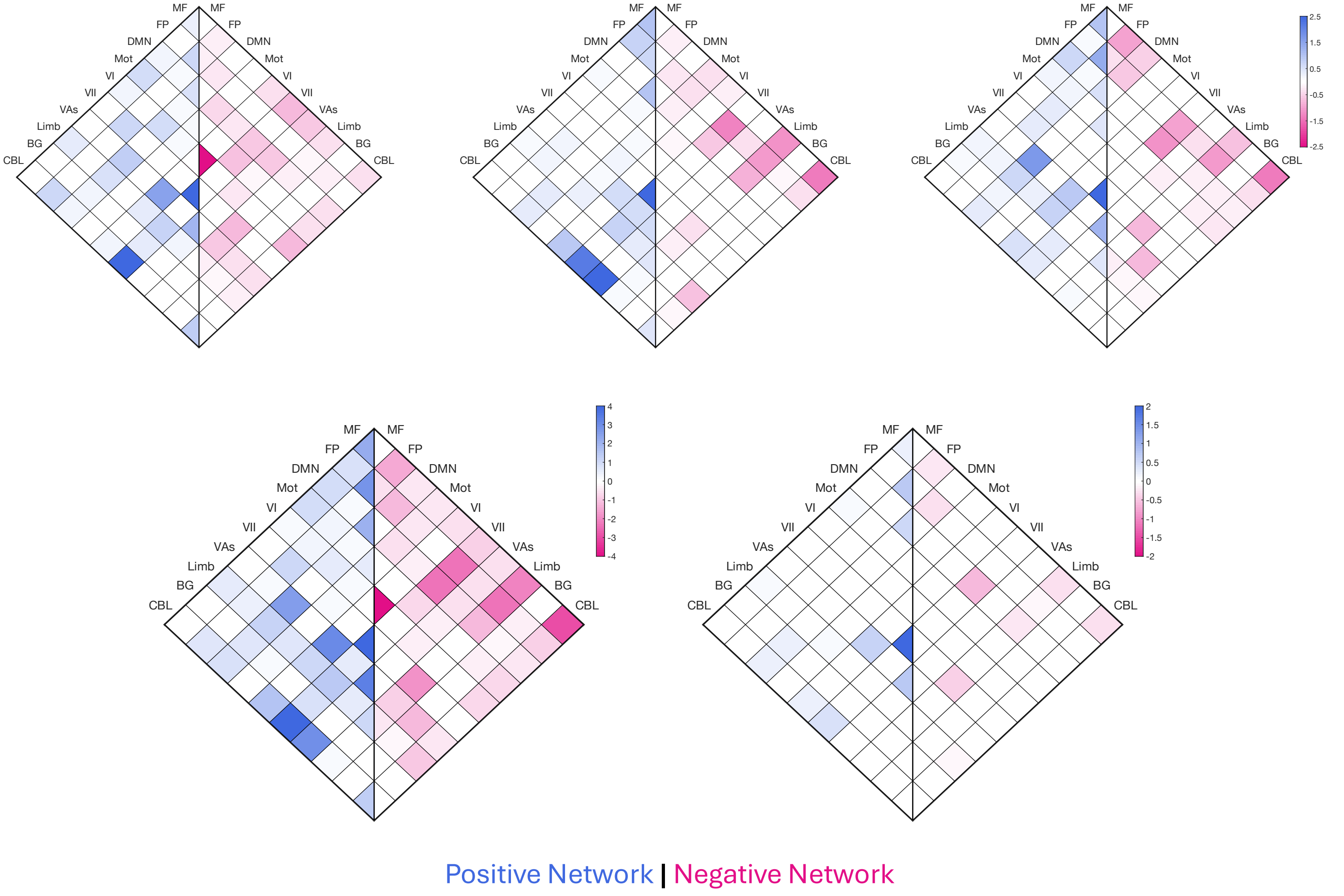


**Supplementary Figure 3.** CPM Canonical network analysis, using resting, volumetric fMRI

Top 3: Flanker, Card Sort, 2-back; Bottom 2: Union, Intersection

Network Acronyms: MF, medial frontal; FP, frontoparietal; DMN, default-mode; Mot, motor; VI, visual A; VII, visual B; VAs, visual association; Limb, limbic; BG, basal ganglia; CBL, cerebellum

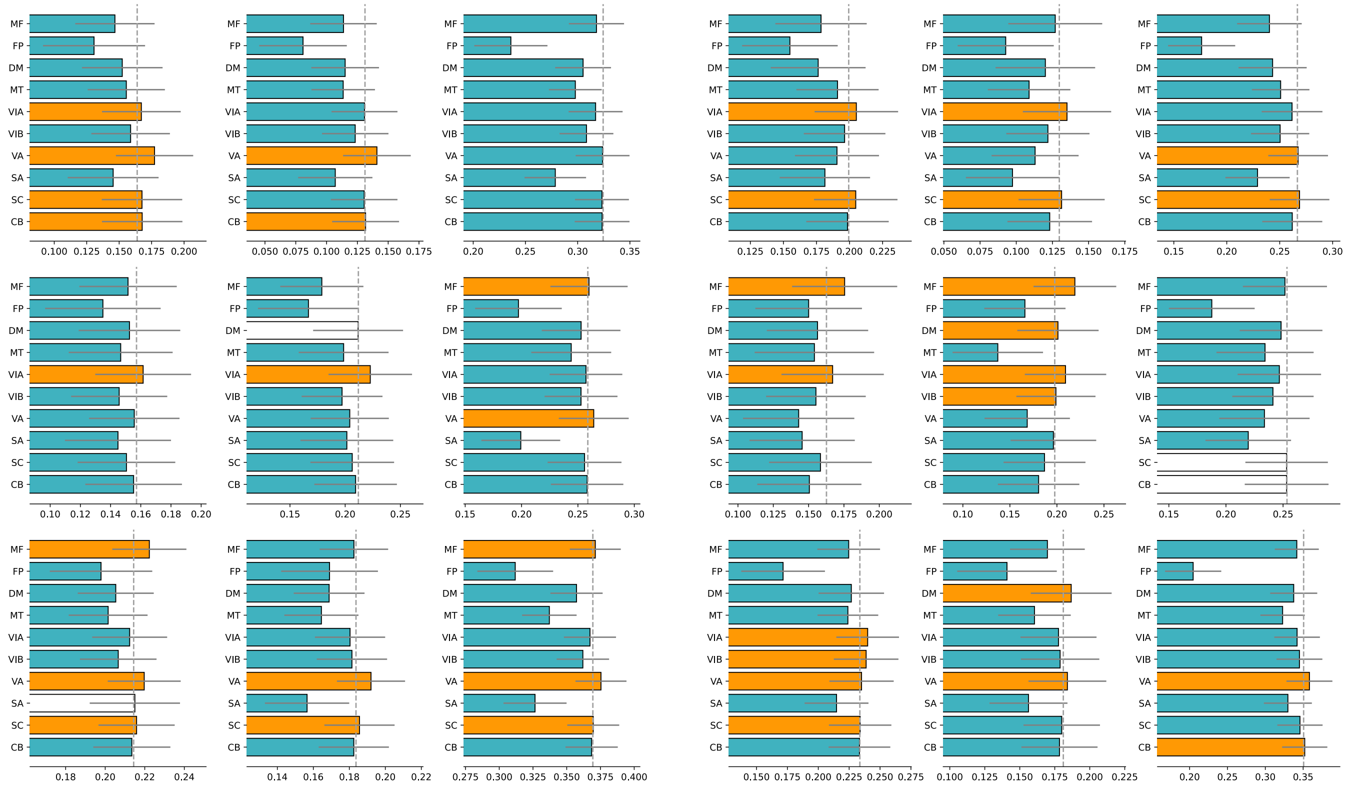

**Supplementary Figure 4.** CPM lesion analysis on 2-Back, volumetric fMRI data.

Left: lesioning positive edge; Right: lesioning negative edges; Rows are training behaviors, in order of: Flanker, Card Sort, 2-back. Columns are testing behaviors, in the same order as rows. Colorings and notations all follow **Figure 2** in the main article. Besides those listed below, all the p-values are < 0.001 after FWE correction.

Exact p-values:

listed in format of (training task-testing task, network name, type of edge, corrected p value)

Flanker-2Back, VA, positive, p=0.007; Card Sort-Card Sort, DM, positive, p=39.91; Card Sort-2Back, MF, positive, p=0.005; 2Back-Flanker, SA, positive, p=0.077; Flanker-2Back, VA, negative, p=0.048; Card Sort-2Back, SC, negative, p=2.89; Card Sort-2Back, CB, negative, p=1.34; 2Back-Flanker, SC, negative, p=0.027;

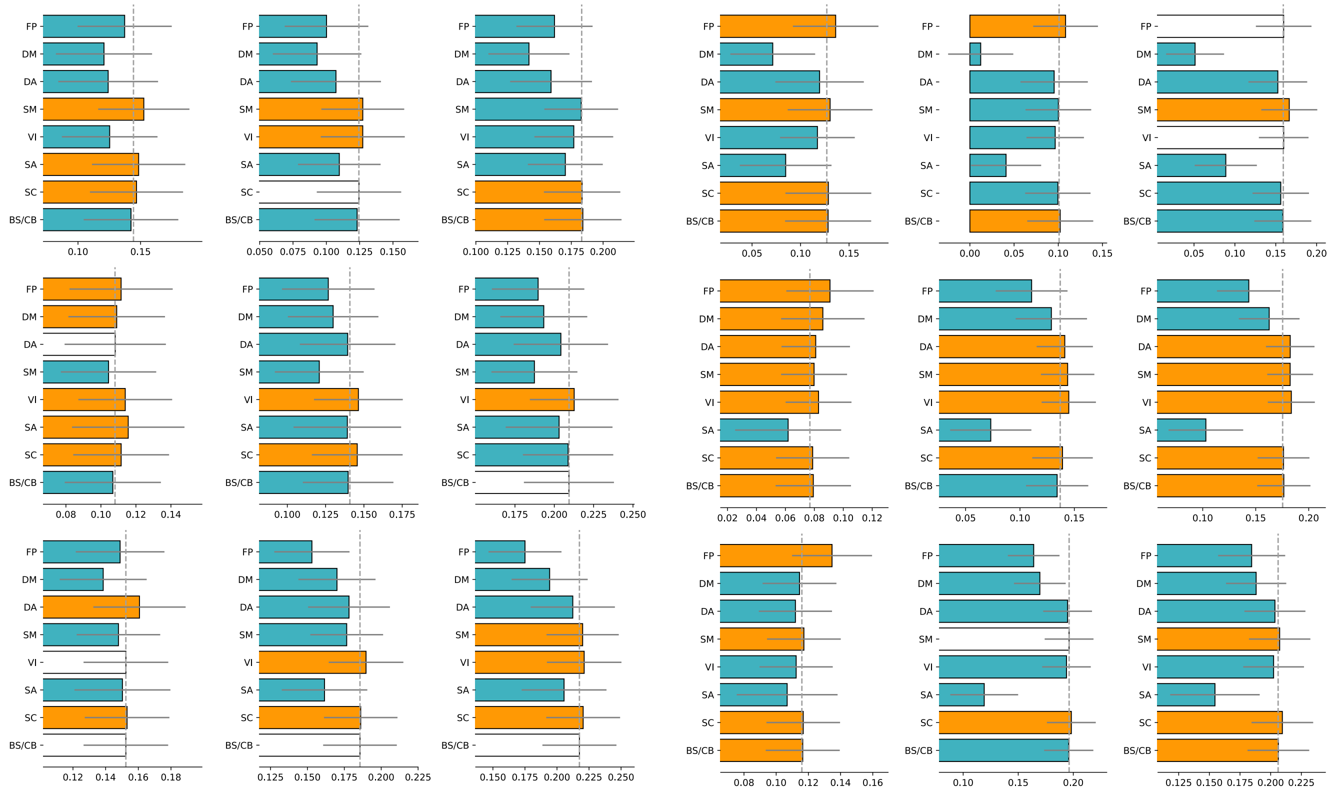

**Supplementary Figure 5.** CPM lesion analysis on resting, grayordinate fMRI data.

Left: lesioning positive edge; Right: lesioning negative edges; Rows are training behaviors, in order of: Flanker, Card Sort, 2-back. Columns are testing behaviors, in the same order as rows. Colorings and notations all follow **Figure 2** in the main article. Besides those listed below, all the p-values are < 0.001 after FWE correction.

Exact p-values:

listed in format of (training task-testing task, network name, type of edge, corrected p value)

Flanker-Card Sort, SC, positive, p=30.34; Flanker-2Back, SM, positive, p=0.020; Card Sort-Flanker, DM, positive, p=0.029; Card Sort-Flanker, DA, positive, p=36.70; Card Sort-2Back, BS/CB, positive, p=0.317; 2Back-Flanker, VI, positive, p=58.67; 2-Back-Flanker, BS/CB, positive, p=Nan; 2Back-Card Sort, BS/CB, positive, p=NaN; 2Back-2Back, BS/CB, positive, p=NaN; Flanker-2Back, FP, negative, p=0.13; Flanker-2Back, VI, negative, p=10.94; 2Back-Card Sort, SM, negative, p=18.53;

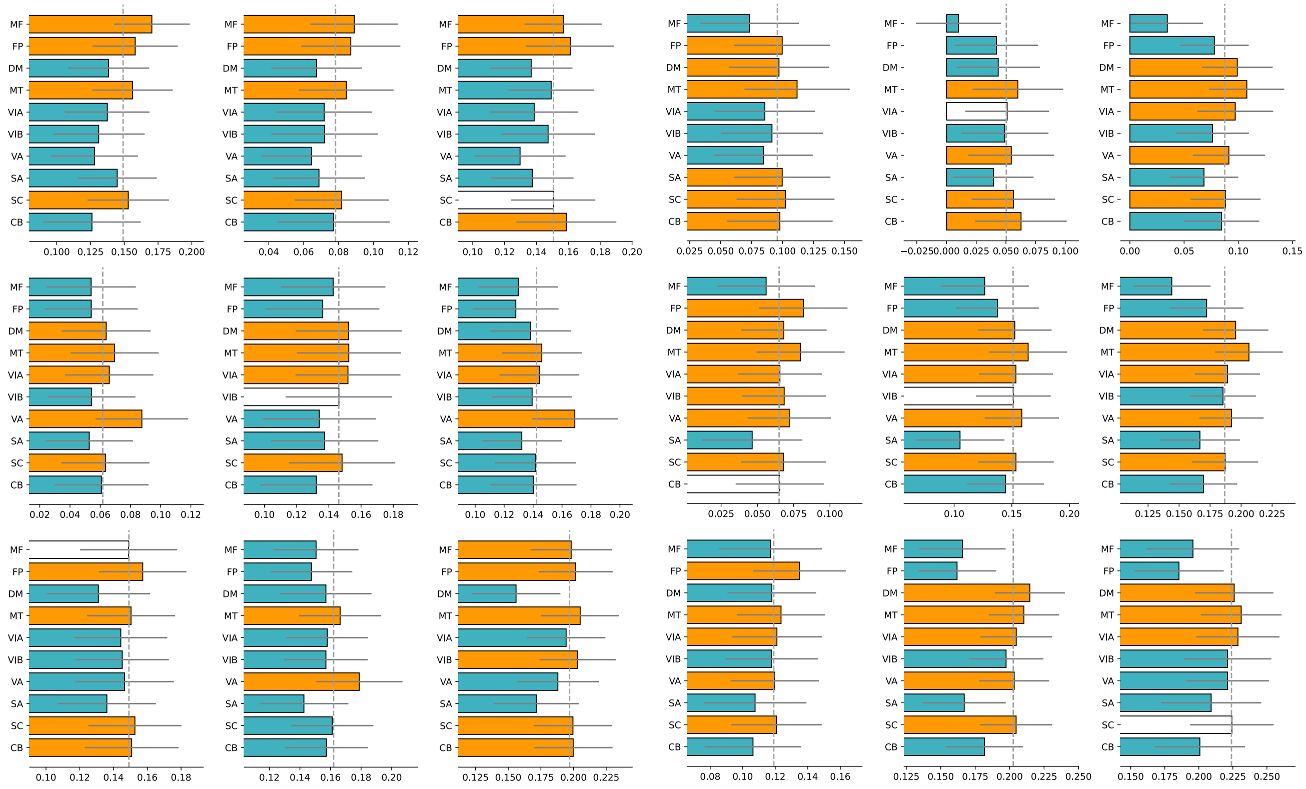


**Supplementary Figure 6.** CPM lesion analysis on resting, volumetric fMRI data.

Left: lesioning positive edge; Right: lesioning negative edges; Rows are training behaviors, in order of: Flanker, Card Sort, 2-back. Columns are testing behaviors, in the same order as rows. Colorings and notations all follow **Figure 2** in the main article. Besides those listed below, all the p-values are < 0.001 after FWE correction.

Exact p-values:

listed in format of (training task-testing task, network name, type of edge, corrected p value)

Flanker-2Back, SC, positive, p=13.87; Card Sort-Flanker, CB, positive, p=0.001; Card Sort-Card Sort, VIB, positive, p=49.90; 2-Back-Flanker, MF, positive, p=27.30; Flanker-Card Sort, VIA, negative, p=0.94; Card Sort-Flanker, CB, negative, p=0.18; Card Sort-Card Sort, VIB, negative, p=13.07; 2Back-2Back, SC, negative, p=0.53.

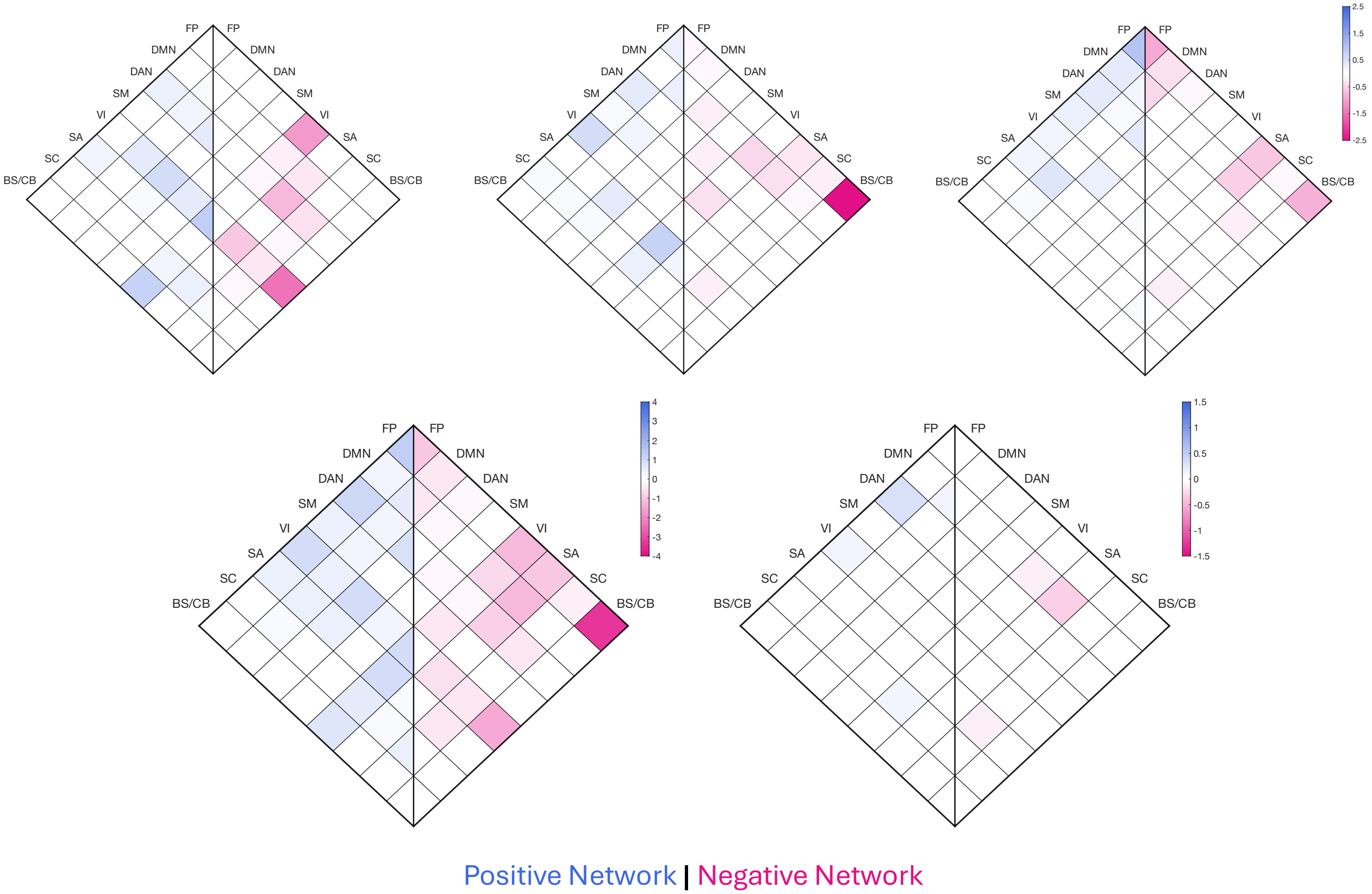


**Supplementary Figure 7.** CPM canonical network analysis of component-specific measures using resting-state, Grayordinate fMRI data.

Top 3: Flanker-specific, Card Sort-specific, 2-back-specific; Bottom 2: Union, Intersection

Network Acronyms: FP, frontoparietal; DM, default mode; DA, dorsal attention; SM, somatomotor; VI, visual; SA, salience; SC, subcortical; BS/CB, brain stem/cerebellum.

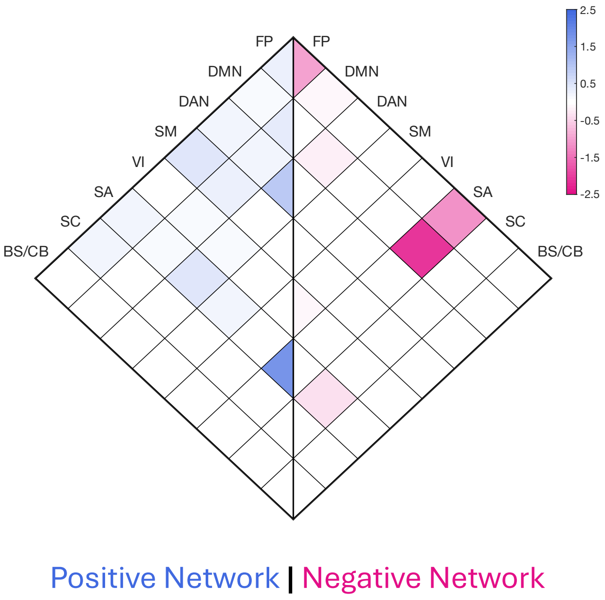


**Supplementary Figure 8.** CPM canonical network analysis of general EF using resting-state, Grayordinate fMRI data.

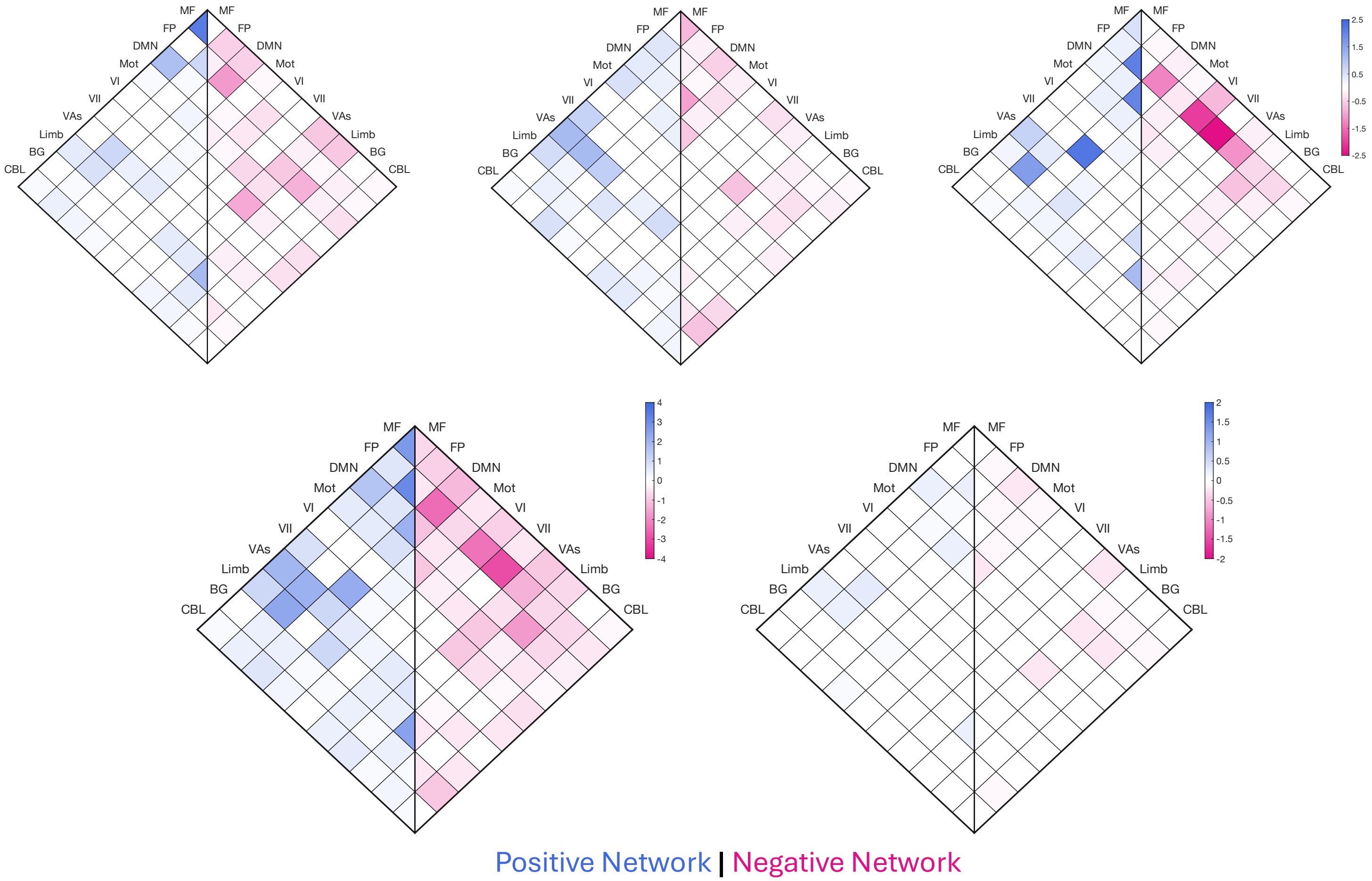


**Supplementary Figure 9.** CPM canonical network analysis of component-specific measures using 2-back, volumetric fMRI data.

Top 3: Flanker-specific, Card Sort-specific, 2-back-specific; Bottom 2: Union, Intersection

Network acronyms: MF, medial frontal; FP, frontoparietal; DMN, default-mode; Mot, motor; VI, visual A; VII, visual B; VAs, visual association; Limb, limbic; BG, basal ganglia; CBL, cerebellum

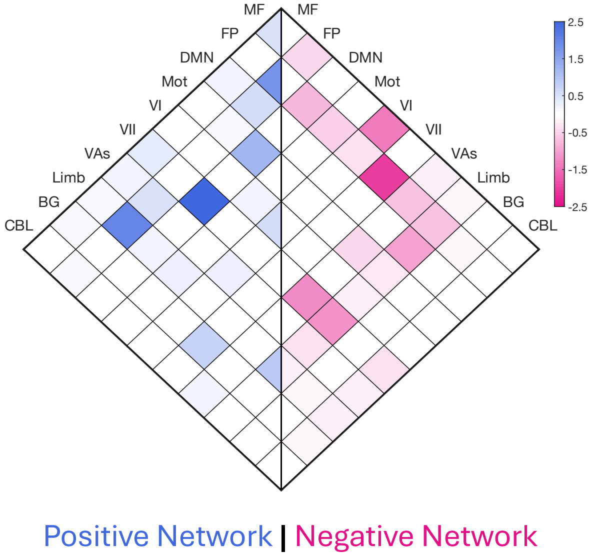


**Supplementary Figure 10.** CPM canonical network analysis of general EF using 2-back, volumetric fMRI data.

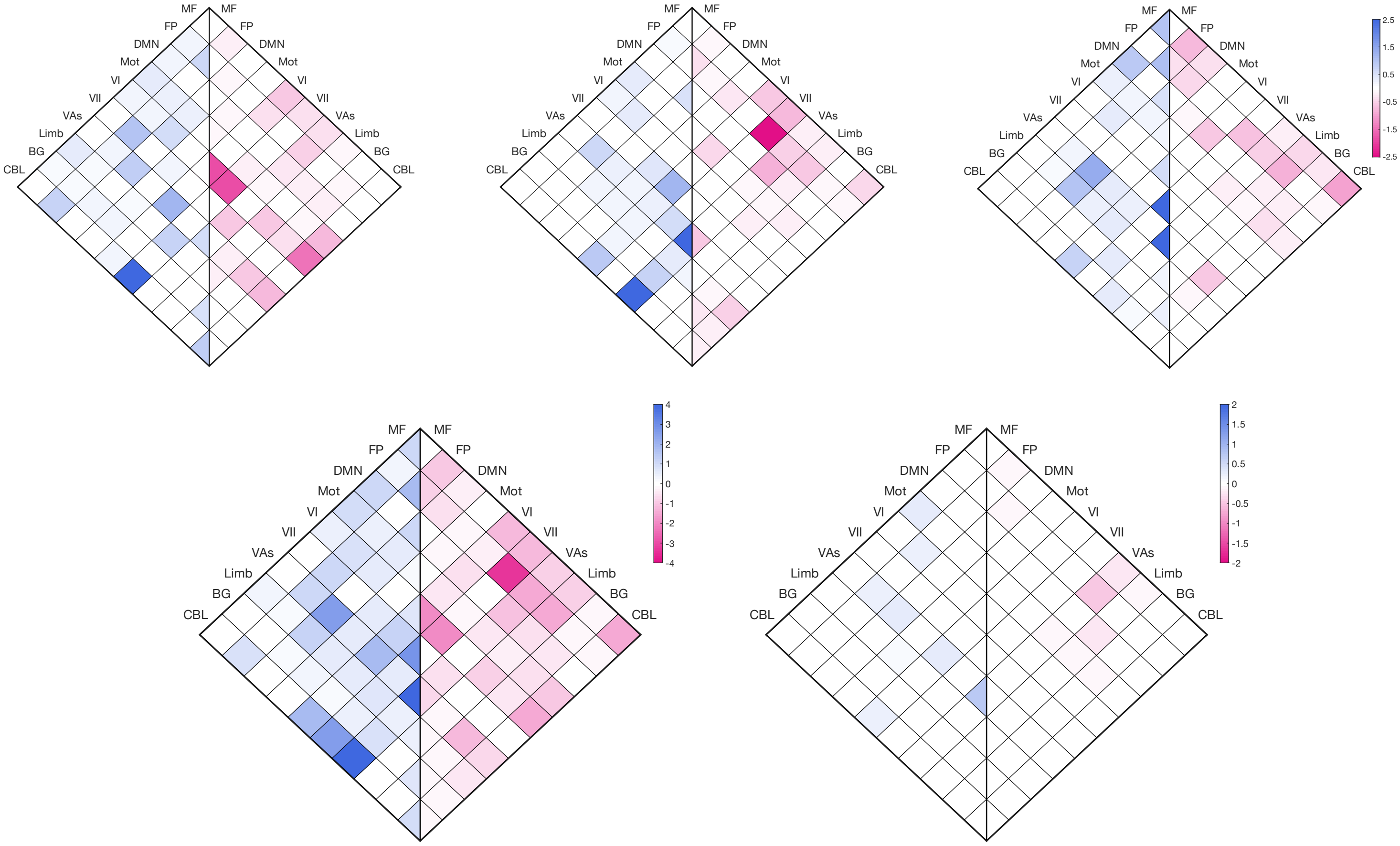


Positive Network | Negative Network

**Supplementary Figure 11.** CPM canonical network analysis of component-specific measures using resting, volumetric fMRI data.

Top 3: Flanker-specific, Card Sort-specific, 2-back-specific; Bottom 2: Union, Intersection

Network acronyms: MF, medial frontal; FP, frontoparietal; DMN, default-mode; Mot, motor; VI, visual A; VII, visual B; VAs, visual association; Limb, limbic; BG, basal ganglia; CBL, cerebellum

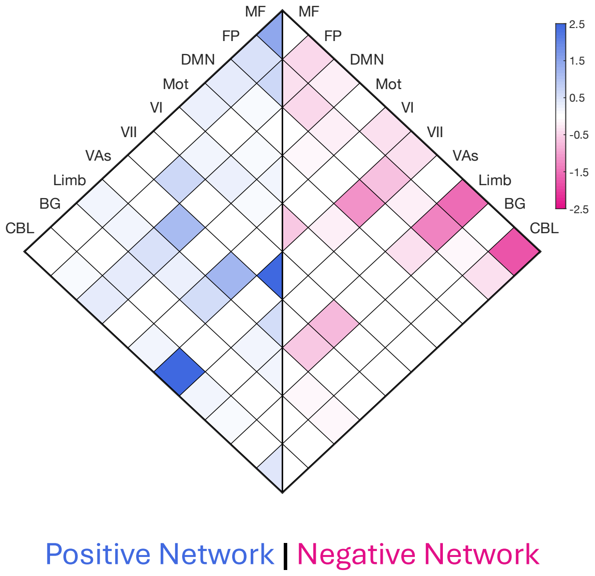


**Supplementary Figure 12.** CPM canonical network analysis of general EF using resting, volumetric fMRI data.
